# Comparing cell penetration of biotherapeutics across human cell lines

**DOI:** 10.1101/2024.03.28.587257

**Authors:** Nefeli Batistatou, Joshua A. Kritzer

## Abstract

A major obstacle in biotherapeutics development is maximizing cell penetration. Ideally, assays would allow for optimization of cell penetration in the cell type of interest early in the drug development process. However, few assays exist to compare cell penetration across different cell types, independent of drug function. In this work, we applied the chloroalkane penetration assay (CAPA) in seven mammalian cell lines as well as primary cells. Careful controls were used to ensure data could be compared across cell lines. We compared the penetration of several peptides and drug-like oligonucleotides and saw significant differences among the cell lines. To help explain these differences, we quantified the relative activities of endocytosis pathways in these cell lines and correlated them with the penetration data. Based on these results, we knocked down clathrin in a cell line with an efficient permeability profile and observed reduced penetration of peptides, but not oligonucleotides. Finally, we used small-molecule endosomal escape enhancers and observed enhancement of cell penetration of some oligonucleotides, but only in some of the cell lines tested. CAPA data provide valuable points of comparison among different cell lines, including primary cells, for evaluating the cell penetration of various classes of peptides and oligonucleotides.

**TOC Graphic:** 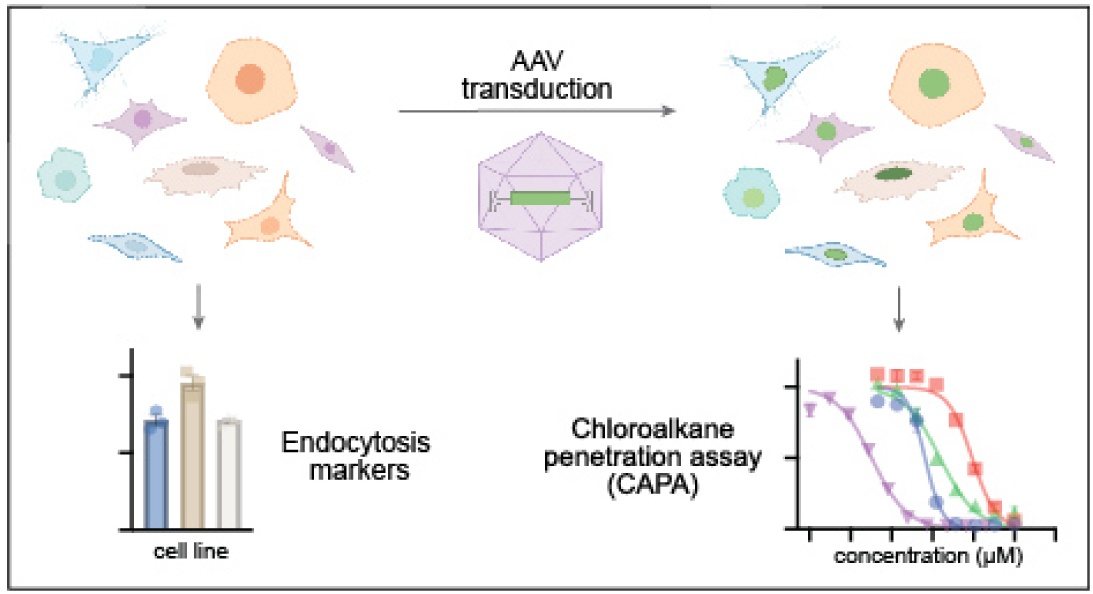

## Introduction

More and more biotherapeutics are entering clinical trials and achieving FDA approval. These include macromolecules across diverse classes including peptides, proteins, siRNAs, and antisense oligonucleotides.^1–3^ Most nucleic acid drugs and many newer peptide drugs have intracellular targets, but historically it has been challenging to maximize the cell penetration of these drug classes. Measuring the cell penetration efficiency of a biotherapeutic in parallel with measuring its cellular activity can be highly beneficial for developing analogs with improved efficacy, and the earlier such information is used to guide development, the better. However, measuring cell penetration of biomolecules is not straightforward. In recent years, chemical biologists have developed a variety of strategies for quantifying passive penetration, total cellular uptake, and cytosolic/nuclear penetration with different detection methods including fluorescence microscopy,^4–7^ mass-spectrometry,^8–13^ NMR,^14^ flow cytometry,^15–17^ FACS,^18,19^ qPCR,^20^ luminescence,^21,22^ or ELISA.^23^ For cell-based assays, the choice of cell line is critical and measurement with different methods and/or in different cell lines can lead to different results.^24–27^ In addition, model cell lines such as HeLa are often convenient, but they do not always recapitulate normal cellular physiology. Further, many drugs require selective delivery to specific cell types that are particularly therapeutically relevant. However, there are few reliable methods to compare cell penetration across different cell lines, particularly for quantitative comparisons and for comparisons of cytosolic/nuclear penetration that are not affected by endosomally trapped material.

Most biotherapeutics including peptides and RNA therapeutics are known to enter mammalian cells by endocytosis.^28–30^ Endocytosis is an energy-dependent process by which compounds are engulfed by the plasma membrane and are initially trapped in vesicles called early endosomes.^31–33^ Early endosomes mature into late endosomes and eventually fuse with lysosomes, where most natural biomolecules are degraded. This process is sometimes termed “non-productive uptake” because material may become associated with the cell but cannot escape endosomes to affect their target(s) in the cytosol and/or nucleus. For some biotherapeutics, small amounts of the trapped material escape endosomes by poorly defined processes, leading to activity in the cytosol and/or nucleus.^30^ This process is sometimes termed “productive uptake.” The roles of different endocytosis pathways in biotherapeutic uptake have been explored, commonly by using chemical inhibitors which provide only moderate pathway specificity and thus often lead to contradictory results.^34–36^ The most well-studied endocytosis pathways are macropinocytosis, clathrin-mediated endocytosis, and caveolin-mediated endocytosis.^33^ Established markers for these pathways include high-molecular-weight dextran for macropinocytosis,^33,37,38^ transferrin for clathrin-mediated endocytosis,^33,39–41^ and cholera toxin subunit B for caveolin-mediated endocytosis.^42,43,34^ Importantly, different cell lines have different basal rates for each of these pathways,^33^ which could partially explain the differences in biomolecule uptake, cytosolic penetration, and activity observed in different cell lines.^27^

We were interested in comparing the relative degrees of cell penetration of peptide and oligonucleotide therapeutics across different cell lines, and we sought to correlate those results to the basal rates of endocytosis in those cell lines. To do so, we employed the Chloroalkane Penetration Assay (CAPA) developed in 2017 in the Kritzer lab.^15,44,45^ CAPA is a high-throughput assay that exclusively reports on the productive uptake of a compound. CAPA relies on HaloTag,^46^ an enzyme that covalently reacts with a small chloroalkane tag. Cells are induced to express HaloTag either in the cytosol or in the nucleus, so that CAPA signal is exclusively attributed to material that accessed the cytosol or nucleus. Compounds-of-interest are prepared with a small chloroalkane tag (ct-compound) and cells are treated with serial dilutions of the ct-compound (*pulse* step). After a specific incubation time, cells are washed and chased with chloroalkane-tagged tetramethylrhodamine (ct-TAMRA) that readily enters cells and reacts with any HaloTag that was unreacted in the pulse step. Finally, unreacted dye is washed out and fluorescence is measured using benchtop flow cytometry. In-plate controls include untreated cells which represent background (0% fluorescence) and cells treated with only ct-TAMRA which represent 100% fluorescence. These controls allow the flow cytometry data to be normalized and fit with a sigmoidal curve that provides a “CP_50_” value, the concentration of ct-compound at which 50% of cytosolic/nuclear HaloTag was blocked in the pulse step. The smaller the CP_50_, the more cell-penetrant the ct-compound. CAPA has been broadly adopted in industry and academia, and it has been successfully applied to peptides,^15,47–50^ antisense oligonucleotides and siRNAs with different lengths and chemical modifications,^45,51,52^ cyclotides,^53^ peptidomimetic macrocycles,^54–56^ PROTACs,^57^ and lipid nanoparticle-mediated delivery of proteins.^58^ It has even been adapted for 3D tissue models to measure deep penetration of biotherapeutics,^59^ and for bacterial cells to evaluate the bacterial penetration of small molecules.^60^

We recently reported the ability to use adeno-associated virus to deliver HaloTag to different cellular compartments and different cell types.^45^ Building on this work, we sought to employ CAPA in many different cell lines, including primary cells, in order to investigate the dependence of cell penetration of peptide and oligonucleotide therapeutics on cell type and basal rates of different endocytosis pathways.

## Results

### CAPA in seven different mammalian cell lines

In order to carry out CAPA in a cell line of interest, a HaloTag expression construct needs to be delivered. For this work, we chose to deliver HaloTag fused to GFP and histone 2B, because this was the most robust adeno-associated virus (AAV) construct in our hands and because nuclear and cytosolic penetration have shown very similar patterns in prior data with peptides and oligonucleotides.^15,45^ We delivered this construct to seven different cell lines using previously described AAV vectors.^45^ The cell lines were chosen to represent different cell types, tissues of origin, and different reported susceptibilities to peptide and/or oligonucleotide therapeutics.^26,27^ After transduction, we observed a shift in green fluorescence compared to non-transduced cells, indicating that the fusion construct was expressed in all seven cell lines (**Figure 1A** and **SI Figure 3**). To obtain broadly comparable levels of HaloTag expression, we empirically optimized transduction conditions, including multiplicity of infection (MOI) and exposure times, for each cell line (**SI Tables 1-2**). For example, U2OS cells required a MOI of 10^3^ and 24 hours of incubation, while HepG2 cells required a MOI of 10^6^ and 48 hours of incubation to produce a comparable GFP expression profile. We next verified that active HaloTag was being produced by labeling cells with ct-TAMRA, similar to the chase step of CAPA. We observed a shift in red fluorescence for transduced cells following incubation with ct-TAMRA compared to transduced, non-labeled cells (**Figure 1B**). Having verified that active HaloTag was present, we then sought to empirically measure the concentration at which HaloTag would get saturated with ct-TAMRA, in order to ensure CAPA experiments used sufficient ct-TAMRA in the chase step (**Figure 1C** and **SI Figure 4**). For this experiment, serial dilutions of ct-TAMRA were added to HaloTag-expressing cells. We defined SAT_50_ as the concentration of ct-TAMRA required to produce half-maximal cellular labeling – the concentration required to saturate 50% of the expressed HaloTag. Despite optimization, there were some differences among the cell lines for their green fluorescence distributions (GFP expression), red fluorescence distributions following ct-TAMRA labeling (active HaloTag levels), and SAT_50_ values (ct-dye required to saturate 50% of the expressed HaloTag). As described below, the differences were small enough that, with additional controls, we concluded that we could reasonably compare CAPA results across these different cell lines.

**Figure 1.**
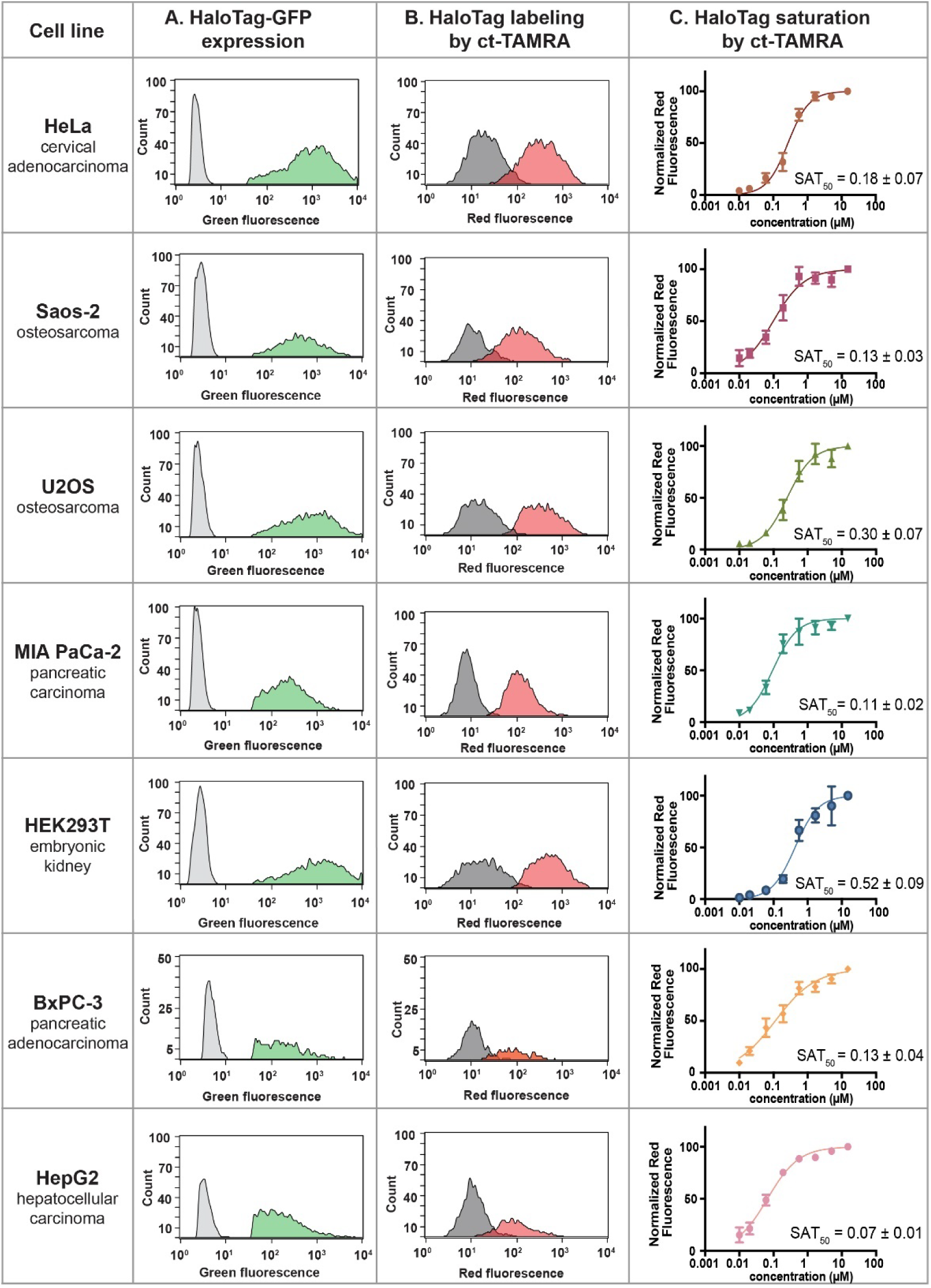
HaloTag expression in seven human cell lines. **(A)** Expression of HaloTag-GFP was measured as a shift in green fluorescence of AAV-transduced cells (green) compared to non-transduced cells (grey). **(B)** Expression of functional HaloTag was measured as a shift in red fluorescence of transduced cells treated with 5 μM ct-TAMRA for 15 min (red) compared to transduced, non-labeled cells (grey). Flow cytometry histograms show representative data from one of at least three independent trials. **(C)** Transduced cells were pulsed with only optiMEM for 4 h or 24 h and then chased with serial dilutions of ct-TAMRA for 15 min. See **SI Figure 4** for similar saturation curves upon 4 h incubation. Data are averages from three independent trials and error bars show standard error of the mean. SAT_50_ values are reported as the average and standard error of the mean from three independent curve fits to three independent trials.

We carried out CAPA experiments for a panel of six compounds (**SI Table 3**): two cell-penetrating peptides (ct-R9W and ct-Tat), a hydrocarbon-stapled peptide (ct-SAHB),^61^ an antisense oligonucleotide with phosphorothioate backbone and 2’-methoxyethyl modified sugars corresponding to the FDA-approved drug nusinersen (ct-nusinersen), a phosphorodiamidate morpholino oligomer (ct-PMO), and a small molecule that commonly used as a control (ct-W; compounds further described in **SI Table 3**). Informed by our prior work applying CAPA to peptide and oligonucleotide therapeutics, we tested two different time points: 4-hour incubations for ct-R9W, ct-Tat, ct-SAHB, and ct-W (**SI Figures 5 and 7**) and 24-hour incubations for ct-nusinersen, ct-PMO, ct-R9W, and ct-W (**SI Figures 6 and 8**). The CP_50_ values of each compound in each of the cell lines are shown in **Figure 2**, and **SI Table 12**. Most CP_50_ values for the same compound across different cell lines differed from each other with statistical significance (**SI Tables 4-11**). Among the cell lines tested, we observed that the pancreatic cancer cell line MIA PaCa-2 was most permissive for cell penetration of both peptide and oligonucleotide therapeutics, and HEK293T cells were least permissive.

**Figure 2.**
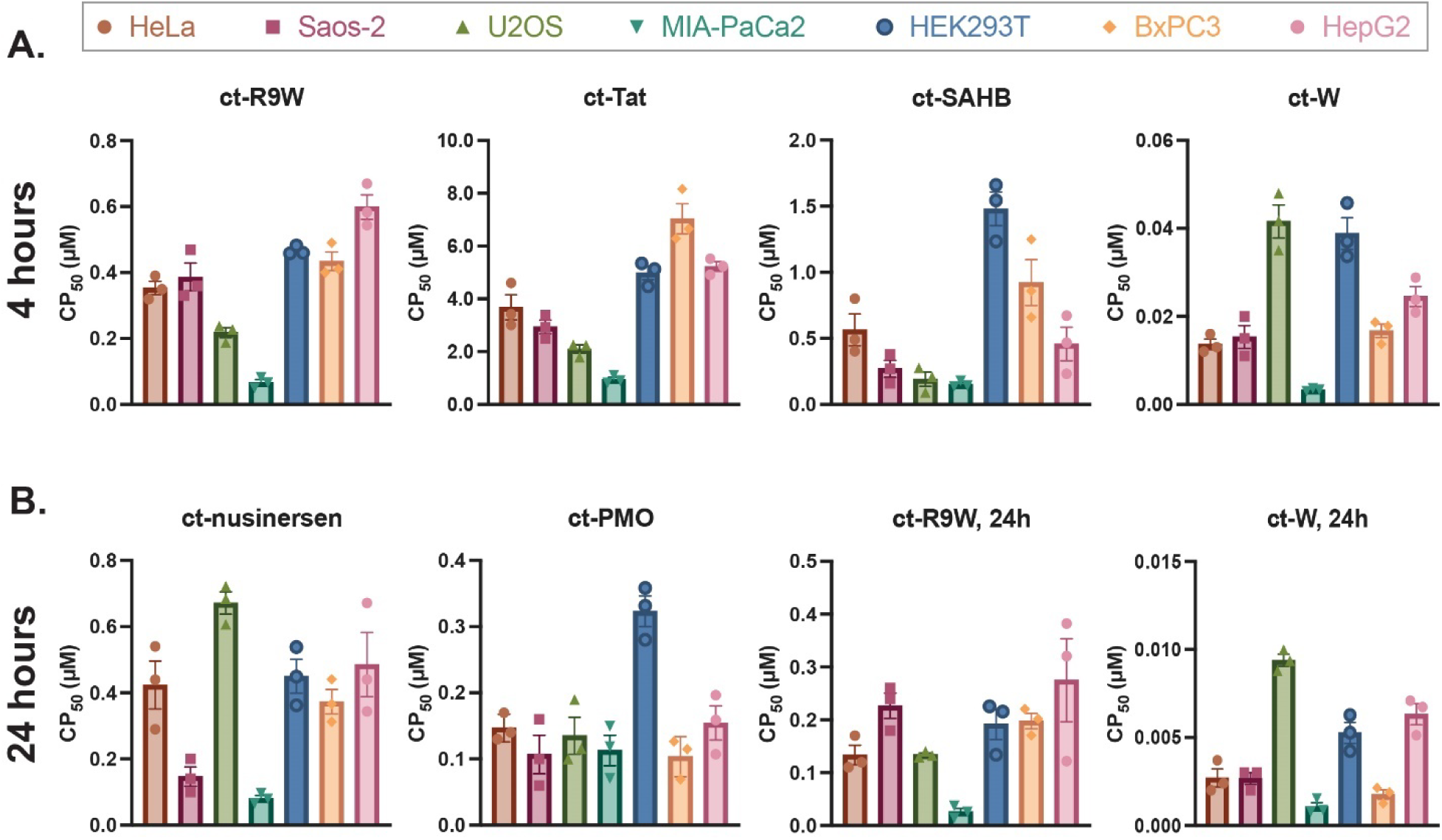
CAPA data for cell-penetrant peptides and oligonucleotides in seven human cell lines. **(A)** CP_50_ values for ct-peptides and controls in seven cell lines after 4 h incubation. These values are derived from raw data and curve fits shown in **SI Figure 5**. **(B)** CP_50_ values for ct-oligonucleotides and controls in seven cell lines after 24 h incubation. These values are derived from raw data and curve fits shown in **SI Figure 6**. CP_50_ values are reported as the average and standard error of the mean from three independent curve fits to three independent trials. Numerical values of CP_50_s are provided in **SI Table 12** and statistical significance tests are provided in **SI Tables 4-11**.

As mentioned above, transduction conditions were optimized for uniform production of HaloTag but there were still some differences among the cell lines (**Figure 1A**, **SI Figure 3**). Having observed different CP_50_ values for the same compound across different cell types, we next sought to investigate the extent to which these values correlated with differences in HaloTag expression. If CP_50_ values and HaloTag expression were not correlated, then the CP_50_ values likely reflected intrinsic properties of those cell lines relevant for drug delivery. First, we compared data from cells gated broadly for HaloTag-GFP expression (the normal gate applied in CAPA, **SI Figure 3, SI Figure 9**, **SI Table 13**)^15,44^ to data from cells gated with a narrower green fluorescence gate (**Figure 2**, **SI Figure 3, SI Table 12**). The narrower gate effectively standardized the range of HaloTag expression across the different cell lines. Importantly, the narrower gate altered the absolute CP_50_ values but they did not change the relative CP_50_ values for any compounds compared across different cell lines. There were also differences in the concentration of ct-dye required to saturate the expressed HaloTag (SAT_50_ values, **Figure 1C**) which depends on HaloTag expression levels but also ct-dye permeability. To further examine whether the CP_50_ values were dependent on HaloTag expression levels, we plotted the CP_50_ values of each ct-compound against the SAT_50_ values for every cell line (**SI Figures 10, 11**) and tested whether there was a linear relationship between them. The plots and the R^2^ values for linear curve fits showed that there were not linear relationships between the CP_50_ and SAT_50_ values. These independent controls indicated that differences among the CP_50_ values across different cell lines were not primarily due to different HaloTag levels in those cell lines.

We were surprised by how different the CP_50_ values were among the cell lines, even for the small-molecule control ct-W which is passively penetrant. Even for ct-W, plots of CP_50_ values against the mean levels of HaloTag-GFP present in each cell line showed no linear relationship (**SI Figure 12**). In fact, there were cell lines with similar CP_50_ for ct-W, but different mean levels of HaloTag (HeLa, Saos-2, BxPC3) and cell lines with similar average levels of HaloTag expression, but different CP_50_ values (MIA PaCa-2, BxPC3, HepG2). The same was true for comparing CP_50_ values and SAT_50_ values for ct-W – no linear relationships were observed, cell lines with similar SAT_50_ values had different CP_50_ values, and cell lines with similar CP_50_ values had different SAT_50_ values (**SI Figures 10, 11**). These comparisons provided further evidence that the different CP_50_ values represent different intrinsic properties of each cell line, such as penetration and distribution of compounds, nonspecific or specific binding to cellular components, cell morphology and surface area, and most importantly different endocytosis activities (see below). While some differences in CP_50_ values across cell lines may arise due to differential expression of HaloTag, these controls support the conclusion that comparing CP_50_ values across cell lines highlights differences in the intrinsic cell penetration properties of those cell lines.

### Correlating CAPA data with endocytosis activities

To account for differences in cell penetration observed among different cell lines, an attractive hypothesis is that the cell lines have different relative endocytosis activities. To correlate cell penetration and endocytosis, we carried out uptake assays in each cell line with known markers for three different endocytosis pathways. Specifically, we treated cells with fluorescently labeled high-molecular-weight dextran, transferrin, and cholera toxin subunit B, which are markers for uptake via macropinocytosis, clathrin-mediated endocytosis, and caveolin-mediated endocytosis, respectively. After 1 hour of incubation at 37°C, cells were washed and fluorescence was measured by flow cytometry and normalized to background to generate values in relative fluorescence units (**SI Table 14**, **SI Figure 13**). Control experiments performed at 4°C showed reduced uptake levels for all three processes, as expected (**SI Figure 13**). There were many differences in the relative activities of each pathway across the seven cell lines (**Figures 3A, 3B**). HEK293T cells had, by far, the highest uptake of all markers – for all three pathways, HEK293T cells had roughly double the uptake compared to the next-most-active cell line. Among the rest of the cell lines studied, Saos-2 and MIA PaCa-2 cells had the highest macropinocytosis activity, U2OS and MIA PaCa-2 cells had the highest clathrin-mediated endocytosis activity, and BxPC3 cells had the highest caveolin-mediated endocytosis activity. HepG2 cells had the lowest macropinocytosis and clathrin-mediated endocytosis activities, and U2OS cells had the lowest caveolin-mediated endocytosis activity.

**Figure 3.**
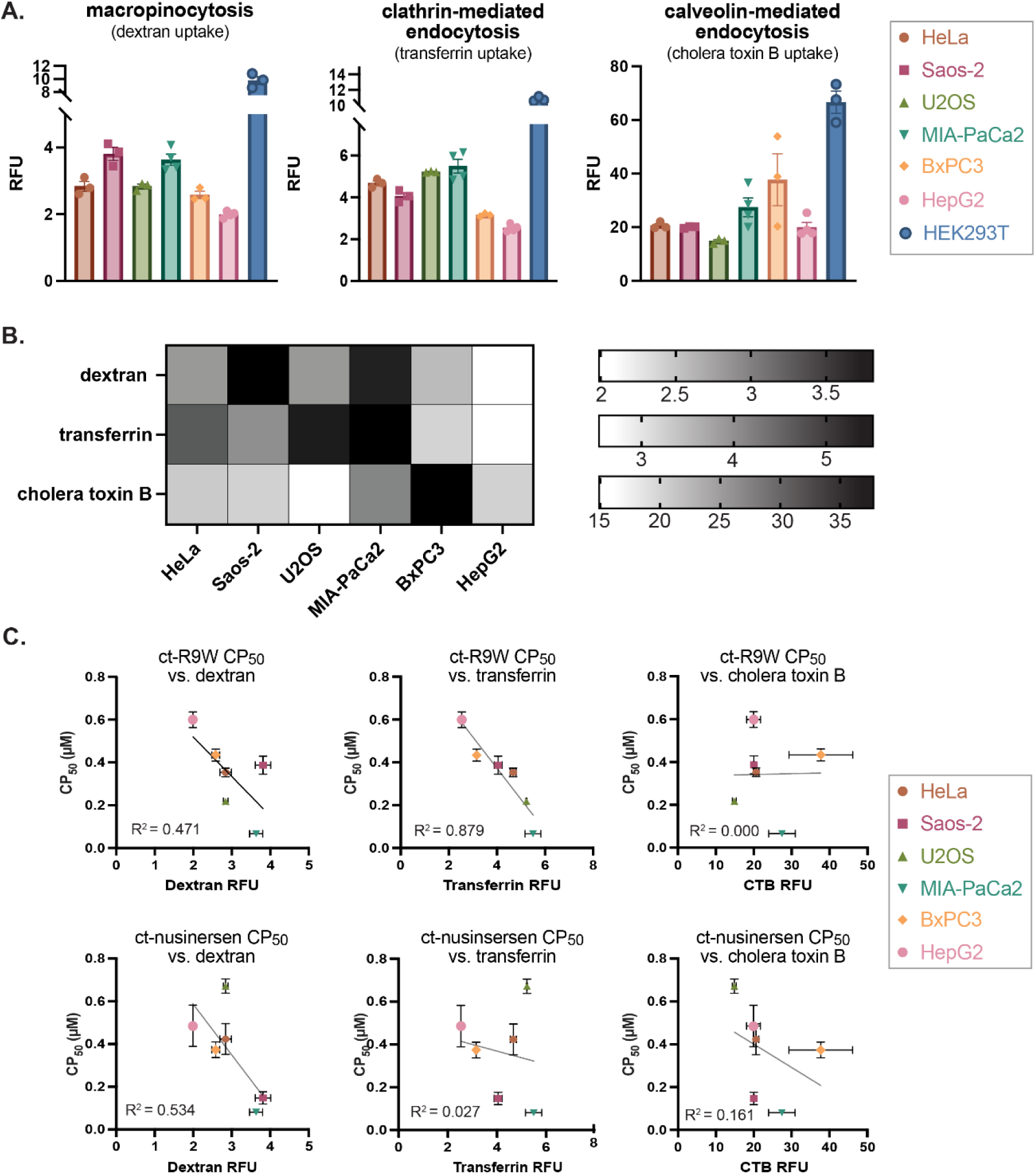
Correlation of cell penetration data with endocytosis activities. **(A)** Relative activities of different endocytosis pathways in different cell lines were measured using 1 mg/mL TAMRA-labeled high-molecular-weight dextran for macropinocytosis, 1 mg/mL TAMRA-labeled transferrin for clathrin-mediated endocytosis, and 5 μg/mL AlexaFluor-647-labeled cholera toxin B for caveolin-mediated endocytosis after 1 hour incubation at 37°C. **(B)** Heat maps showing relative activities of endocytosis pathways in six cell lines (HEK293T cells had very high activity in all three pathways and were considered separately). **(C)** Plots examining possible correlations between cell penetration (CP_50_, lower values indicate greater cell penetration) and endocytosis activities in six of the cell lines for ct-R9W and ct-nusinersen. Remaining plots are shown in **SI Figure 14**. RFU and CP_50_ values are reported using the average and standard error of the mean from three independent trials.

The relative endocytosis activities in the cell lines appeared to explain some, but not all, of the CAPA results. For example, the greatest penetration (lowest CP_50_ values) were observed in MIA PaCa-2 cells for all peptides and for ct-nusinersen, and greater penetration was also observed in U2OS cells for the peptides and in Saos-2 cells for ct-nusinersen (**Figures 2B, 2C**). These were also the cell lines with the highest macropinocytosis and clathrin-mediated endocytosis activities (**Figures 3A, 3B**). Similarly, the HepG2 and BxPC3 cell lines had poorer penetration overall (higher CP_50_ values) and these were also the cell lines with less macropinocytosis and clathrin-mediated endocytosis activities. Interestingly, HEK293T cells had the highest endocytosis activities for all three pathways, yet had among the poorest penetration (highest CP_50_ values) of all the cell lines tested. CAPA reports on cytosolic/nuclear penetration, so high CP_50_ values in a cell line with high endocytosis activity may suggest an unusually low degree of endosomal escape. Because of these outlying features, and the common observance of both adherent and suspended cells in HEK293T cultures, we omitted HEK293T cells from subsequent analyses.

To look for global correlations between individual endocytosis pathway activities and cell penetration, we plotted the CP_50_ values for each ct-compound versus the relative activity measurements for individual endocytosis pathways in each cell line (**Figure 3C** and **SI Figure 14**). There were significant correlations for some of the ct-compounds with specific endocytosis pathways. Specifically, the polycationic cell-penetrating peptides ct-R9W and ct-Tat had linear correlations with clathrin-mediated endocytosis activity, implying that cell penetration directly depends on uptake via this pathway. A similar correlation was observed for ct-nusinersen and macropinocytosis activity (**Figure 3C** and **SI Figure 14**), and this correlation was even stronger if the U2OS cell line was omitted from the analysis. For the ct-PMO, there was a slight correlation with caveolin-mediated endocytosis activity (**SI Figure 14**). For the rest of the correlation plots for other ct-compounds and endocytosis activities, there were no linear relationships observed between endocytosis activities and cell penetration.

### Effects of endosomal escape enhancers

Some prior work has shown that chloroquine can enhance the endosomal escape of peptides^62^ and oligonucleotides.^63,64^ So, we next examined the effects of chloroquine on cell penetration in BxPC3 cells, one of the least permissive cell lines tested. Unexpectedly, co-incubation of ct-compounds with 60 μM chloroquine in BxPC3 cells did not increase cell penetration for any peptides or oligonucleotides tested (**SI Figures 15 &16, SI Table 15**). Most compounds tested, including ct-W, showed a two-to-three-fold increase in CP_50_ value upon chloroquine treatment, indicating poorer cell penetration. Interestingly, the CP_50_ of the stapled peptide ct-SAHB increased from 0.88 ± 0.20 μM to 6.44 ± 0.17 μM in the presence of chloroquine, indicating a mechanism of cell penetration that is particularly sensitive to alterations in lysosomal function (**SI Figure 15A**).

Several recent papers have reported compounds that more selectively enhance endosomal escape, especially for antisense oligonucleotides. Such compounds have been called oligonucleotide enhancer compounds (OECs). We chose OECs UNC10217938A^6,65^ and SH-BC-893^66^ to examine their effects on cell penetration of ct-nusinersen and ct-PMO in BxPC3 and U2OS cells. We chose these two cell lines because they were among the cell lines with higher CP_50_ values, and we wanted to examine whether OECs could improve cell penetration as measured by CAPA.

In BxPC3 cells, co-incubation of ct-nusinersen with 10 μM UNC10217938A or 5 μM SH-BC-893 for 24h decreased the CP_50_ value by roughly 30% (**Figure 4A**, **SI Table 16**, **SI Figure 17**). In U2OS cells, 10 μM UNC10217938A decreased the CP_50_ value for ct-nusinersen by roughly 66% (from 0.50 ± 0.02 μΜ to 0.17 ± 0.08 μM) but 5 μM SH-BC-893 had no effect (**Figure 4B**, **SI Table 17, SI Figure 19**). Neither OEC had a significant effect on the measured CP_50_ value for ct-PMO or the small molecule control ct-W (**SI Figure 18B**). These data suggest that OECs can enhance cell penetration of drug-like oligonucleotides, but moderately and in a compound- and cell-specific manner.

**Figure 4.**
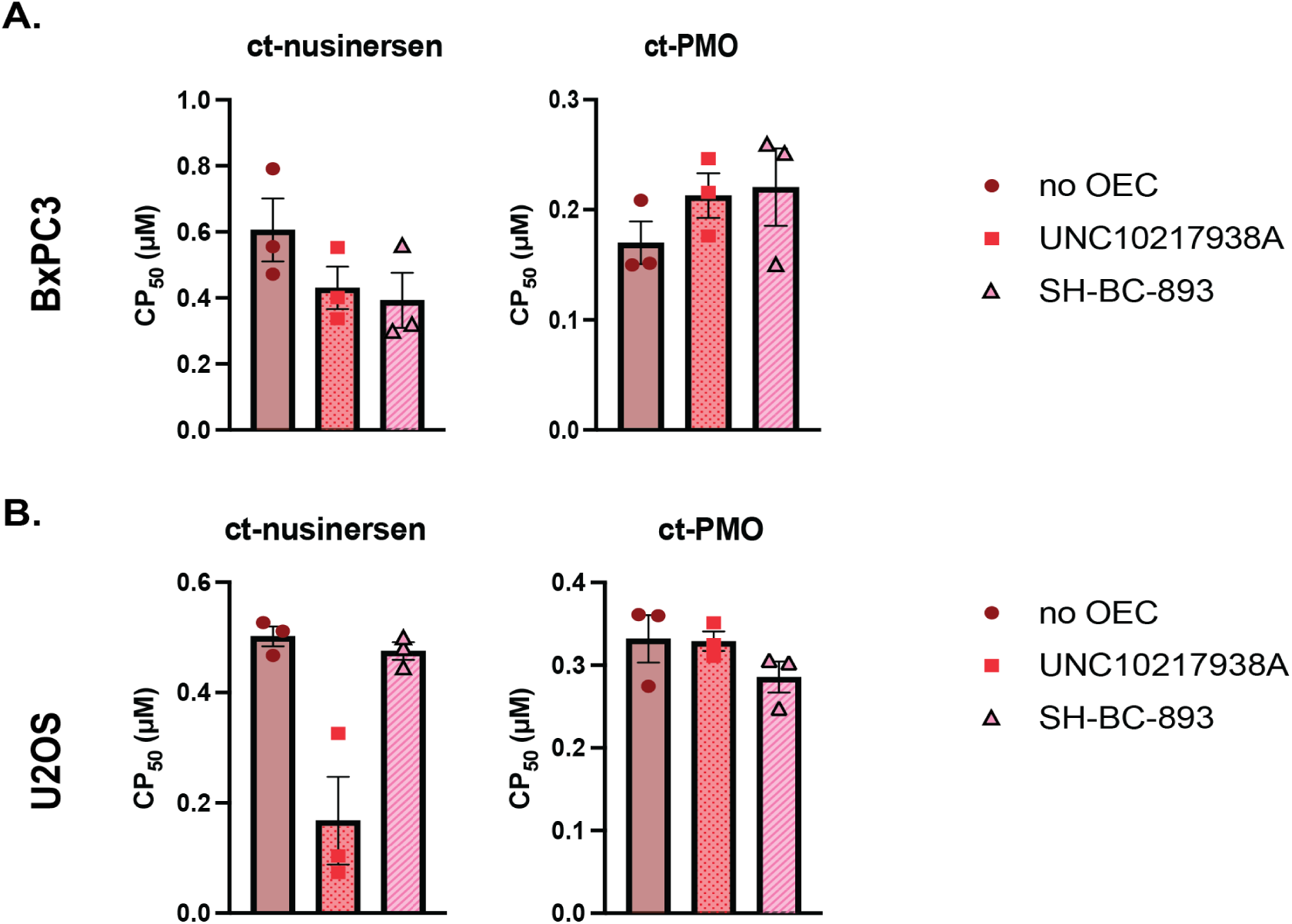
Effects of oligonucleotide enhancer compounds (OECs) on cell penetration. Cell penetration of ct-nusinersen and ct-PMO was measured in the absence or presence of OECs UNC10217938A (10 μM) and SH-BC-893 (5 μM). Selected data are shown here for key comparisons in BxPC3 cells **(A)** and in U2OS cells **(B)**. CP_50_ values are derived from raw data and curve fits shown in **SI Figures 17, 19**. CP_50_ values are reported as the average and standard error of the mean from three independent curve fits to three independent trials. Significance testing for these data are found in **SI Tables 18-21**.

### Effects of clathrin knockdown

Prior work has commonly used endocytosis inhibitors, including amiloride, rottlerin, chlorpromazine, dynasore, methyl-β-cyclodextrin, and nystatin, to attempt to selectively block specific endocytosis pathways. However, these compounds are notoriously non-specific.^33,34,67^ We sought a more specific means of inhibiting a particular pathway, to further examine the relationships between specific endocytosis pathways and cell penetration as measured by CAPA. We chose to knock down clathrin using small interfering RNA (siRNA), which was shown to be a much more robust and selective approach.^34^ We carried out these experiments in MIA PaCa-2 cells since they had the highest levels of clathrin-mediated endocytosis and the lowest CP_50_ values (greatest cell penetration) in general. Transfection conditions and clathrin knockdown were optimized and verified using Western blot (**SI Figure 20**). Upon clathrin knockdown, there was a decrease in the uptake of transferrin, the clathrin-mediated endocytosis maker. The uptake of dextran was not affected much, while the caveolin-mediated pathway seemed to get up-regulated as seen by the significant increase in the uptake of cholera toxin subunit B (**Figure 5B**).

**Figure 5.**
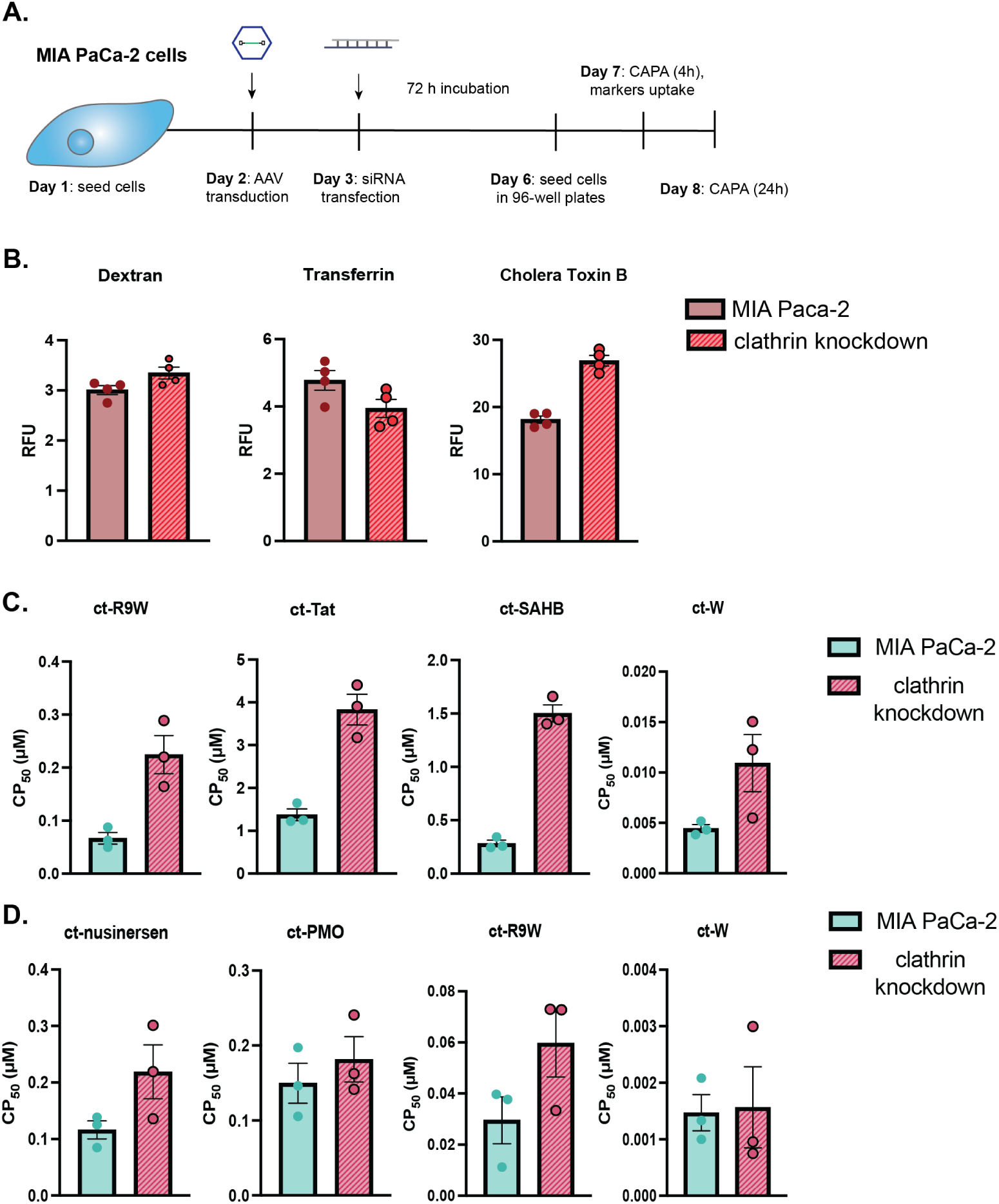
Measuring the effects of clathrin knockdown on cell penetration of ct-compounds in MIA PaCa-2 cells. **(A)** Schematic of experimental plan. **(B)** Effect of clathrin knockdown on uptake of markers of different endocytosis pathways. RFU values are reported as the average and standard error of the mean from three independent trials. **(C)** CP_50_ values for the indicated ct-compounds upon clathrin knockdown after 4 hours of incubation. **(D)** CP_50_ values for the indicated ct-compounds upon clathrin knockdown after 24 hours of incubation. CP_50_ values are reported as the average and standard error of the mean from three independent curve fits to three independent trials, shown in **SI Figure 21**. Numerical CP_50_ values are shown in **SI Table 24**. An unpaired t-test was used (**SI Table 25**, **26**) to calculate *p* values for data in panels B-D.

Cell penetration of the ct-peptides was impaired significantly in the absence of clathrin (**Figure 5C**, **SI Figure 21A**) after 4 hours of incubation. The biggest difference was seen for hydrocarbon-stapled peptide ct-SAHB, whose CP_50_ increased by more than 5-fold (**SI Table 24**), while the penetration of ct-W was not significantly different. After 24 hours of incubation in MIA PaCa-2 cells with clathrin knockdown, the CP_50_ values of the ct-oligonucleotides and ct-R9W appeared to increase, but with a broad spread of values among replicates (**Figure 5D**, **SI Figure 21B**, **SI Table 24**).

### CAPA in primary cells

After comparing cell penetration in seven cultured cell lines using CAPA, we hypothesized that AAV introduction of HaloTag would also allow us to measure cell penetration in primary cells. We used the methods described above to measure endocytosis activities in human umbilical vein epithelial cells (HUVECs, **SI Figure 22**) and we transduced them with AAV (**SI Figure 23**) to enable CAPA measurements for a panel of ct-compounds (**Figure 6A, 6B**). The CAPA data revealed that HUVECs are similarly permissive as many cultured cell lines in terms of cell penetration of peptide and oligonucleotide therapeutics. We were interested whether the data from HUVEC cells would match the correlations observed with cultured cell lines. Penetration of ct-R9W and ct-Tat correlated with transferrin uptake activity (**SI Figure 14**), as observed in cultured cells. However, penetration of ct-nusinersen in HUVEC cells did not correlate with dextran uptake, contrasting with results from most of the cultured cells. We also tested the effects of OEC UNC10217938A on ct-nusinersen penetration in HUVEC cells (**Figure 6C**, **SI Table 27**). UNC01217938A improved the CP_50_ of ct-nusinersen by over 50% in this primary cell line (**SI Figure 24**, **SI Table 27**).

**Figure 6.**
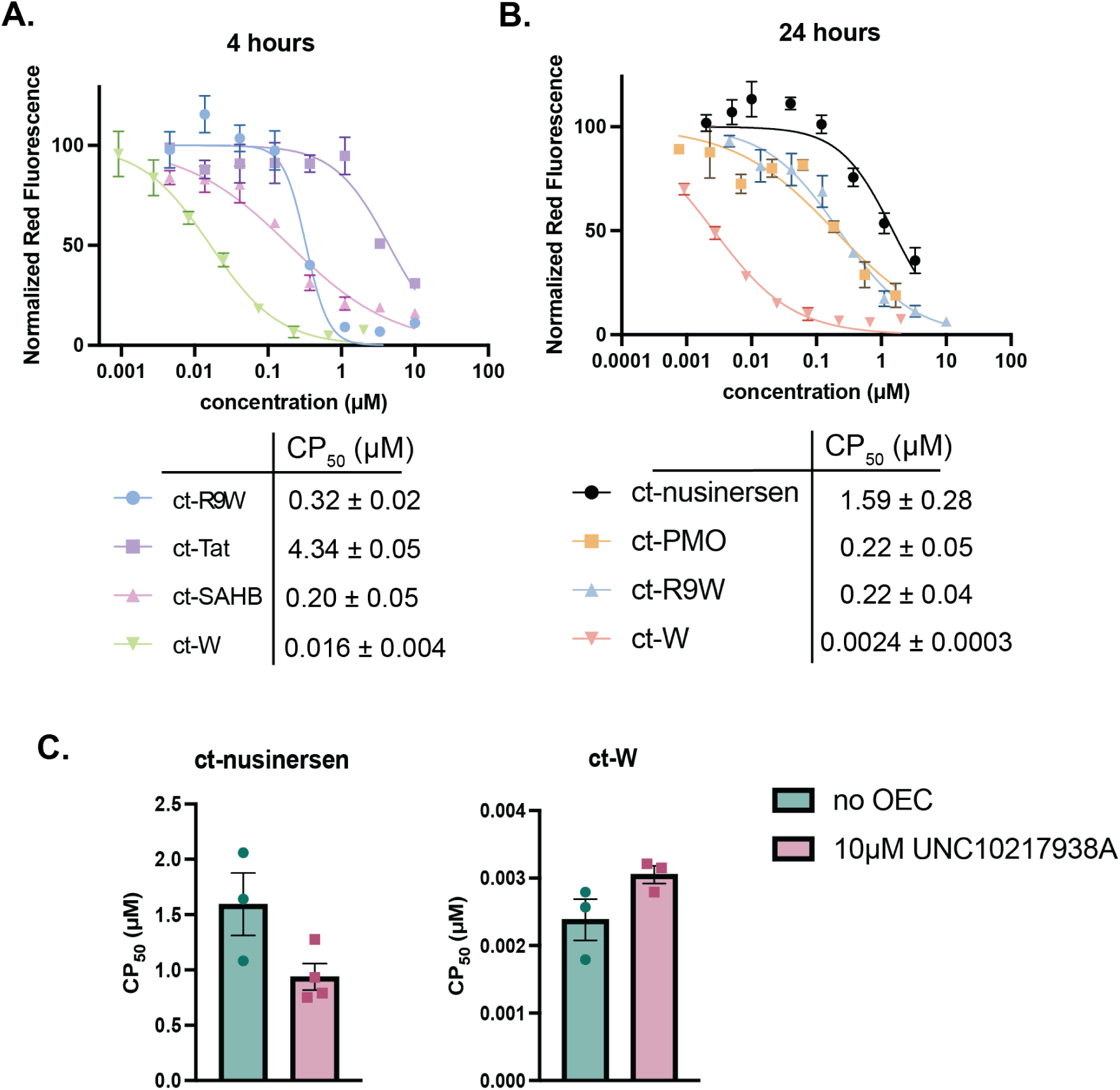
CAPA in primary cells. **(A)** CP_50_ values for the indicated ct-compounds in HUVECs after 4 hours of incubation. **(B)** CP_50_ values for the indicated ct-compounds in HUVECs after 24 hours of incubation. **(C)** CP_50_ values for ct-nusinersen and ct-W in the absence or presence of 10 μM UNC10217938A. All data show results from three independent trials and error bars show standard error of the mean. CP_50_ values are reported as the average and standard error of the mean from three independent curve fits to three independent trials. CP_50_ values in panel C were derived from curve fits shown in **SI Figure 24**.

## Discussion

Evaluating cell penetration in different cell types is crucial for the development of more effective biotherapeutics. The compounds in our panel included cationic cell-penetrating peptides, a hydrocarbon-stapled peptide, and two classes of synthetic oligonucleotides. All compounds showed differences in cell penetration in different cell lines, which was observed in prior literature.^24–27^ For example, studies showed that the total uptake of fluorescently-labeled CPPs^24^ or fluorescently-labeled peptides and peptidomimetics^27^ was different when measured in different cell lines. Similar results were observed for the CPP-mediated delivery of GFP in different cell lines.^25^ Linnane et al. showed differences among cell lines in the total uptake of a label-free ASO by immunofluorescence using an anti-oligonucleotide antibody.^26^ Our results add substantially to these prior works because CAPA measures *productive uptake* of biomolecules, in this case material that reached the nucleus of cells, rather than total cellular uptake.

Among the cell lines tested, we found the pancreatic cancer cell line MIA PaCa-2 to be among the most permissive cell lines for compound penetration, while the pancreatic cancer cell line BxPC3 was one of the least permissive cell lines (**Figure 2**). This can be explained partially by the fact that MIA PaCa-2 cells have an activating *KRas* mutation while BxPC3 cells have wild-type *KRas*, which was observed by others to enhance endocytosis and cell penetration of externally added compounds.^26,27,68^

Our data add to the ongoing examination of the mechanisms of cell penetration for cationic cell-penetrating peptides (such as Tat and polyarginine, both tested in this work).^69^ Such peptides are well-documented to enter cells by endocytosis,^70–76^ but different pathways of endocytosis have been proposed to be responsible for their initial uptake. Evidence has documented that they can be taken up by macropinocytosis,^77–80^ clathrin-mediated endocytosis,^79,81,82^ or caveolin-mediated endocytosis,^83–85^ or all three pathways.^86^ Direct translocation across the membrane has also been proposed, and appears to occur most noticeably at higher peptide concentrations.^5,87–90^ In this work, we obtained data across six different cell lines which show a linear correlation between clathrin-mediated endocytosis activity and cell penetration of ct-R9W and ct-Tat (**Figure 3C**). We observed a similar trend for macropinocytosis activity, especially for ct-R9W in all cell lines except Saos-2 (**Figure 3C**). No relationship was observed between the levels of caveolin-mediated endocytosis and cell penetration of ct-R9W and ct-Tat, and thus our data agree with studies that rule out caveolin-mediated endocytosis as a relevant uptake pathway for cationic peptides.^81^ The notion that clathrin-mediated endocytosis is a primary uptake route for CPPs is also supported by our CAPA data upon clathrin knockdown in MIA PaCa-2 cells (**Figure 5C**), where the CP_50_ values of ct-Tat and ct-R9W increased by ~3-fold.

Hydrocarbon-stapled peptides like ct-SAHB were reported to enter Jurkat T cells by macropinocytosis.^61,90–92^ Our data do not show a direct correlation between cell penetration of ct-SAHB and macropinocytosis activity in the cell lines tested. In fact, we do not see a linear correlation with any of the endocytosis pathways studied, although knockdown of clathrin in MIA PaCa-2 cells reduced cell penetration of ct-SAHB by more than 5-fold (**Figure 5C**). Taken with prior work, our data indicate that the primary uptake route for hydrocarbon-stapled peptides may vary among different cell types.

Our data also add to the ongoing effort to understand cell penetration of chemically modified oligonucleotide therapeutics.^93–95^ Antisense oligonucleotides (ASOs) have been shown to enter cells by endocytosis – more specifically, phosphorothioate ASOs have been reported to enter cells via macropinocytosis, clathrin-mediated endocytosis, or caveolin-mediated endocytosis.^95–97^ Some work has shown that phosphorothioate ASOs interact with cell-surface proteins including epidermal growth factor receptor (EGFR),^98^ which is known to stimulate macropinocytosis,^99^ but other work showed they also can interact with stabilin receptors that internalize them via clathrin-coated pits.^100,101^ Our data show a direct relationship between higher macropinocytosis levels and higher cell penetration efficiency for the phosphorothioate ASO ct-nusinersen across most cell lines tested (**Figure 3C**). We did not observe correlations between cell penetration of ct-nusinersen and clathrin-mediated endocytosis activity or caveolin-mediated endocytosis activity. This is also supported by our clathrin knockdown experiment in MIA PaCa-2 cells, where the CP_50_ of ct-nusinersen did not change significantly upon clathrin knockdown. This also reinforces the notion that the different pathways work in synergy, and they complement each other, especially when one is unavailable i.e. clathrin knockdown.^95^ Our analysis from this work agrees with the findings of Koller *et al,*^102^ who showed that clathrin knockdown did not alter functional activity of ASOs.^102^ For the phosphorodiamidate morpholino oligomer (PMO), we only observed a slight correlation with caveolin-mediated endocytosis activity; this agrees with some prior results as well.^103^ The different observations between the PS-ASO and the PMO highlight that the primary mechanisms of entry for different chemically modified oligonucleotides are likely different.

We excluded HEK293T cells from the cross-cell-line analysis because they appeared to have unusually high levels of endocytosis, yet also had poor cell penetration (high CP_50_ values). These data suggest that although HEK293T cells have more rapid uptake than many other cell lines, they may have slow endosomal escape rates – this finding matches some prior work in this cell line.^104^ Moreover, our CAPA results in HEK293T cells also agree with the conclusions drawn by Arora and coworkers using uptake assays based on dye fluorescence.^27^ That study found that the total uptake of several classes of cell-penetrant peptides, including Tat, was similarly poor in HEK293T and BxPC3 cells (both expressing wild-type *Ras*). Our data focusing on the endosomal escape efficiency agree with these observations, as the CP_50_ values for ct-Tat and ct-R9W are highest in BxPC3 and HEK293T cells, while for ct-SAHB the highest CP_50_ is observed in HEK293T cells.

Clathrin knockdown in MIA PaCa-2 cells decreased the uptake of transferrin, as seen by others.^105,106^ Recently, Itagaki *et al.* noted an increase, rather than a decrease, in the uptake of fluorescently-labeled transferrin when they performed a clathrin heavy chain knockdown in HeLa cells.^34^ During the same experiment, they saw no effect in the uptake of dextran, with matches our observations in MIA PaCa-2 cells, and they also saw a decrease in the uptake of cholera toxin B in HeLa while we observed an increase in MIA PaCa-2. These comparisons highlight that even the effects of direct perturbations such as clathrin knockdown can vary depending on the cell line used.

Discrepancies between uptake activities and cell penetration are typically ascribed to efficiency of endosomal escape. We sought to enhance escape efficiency in one of the least permissive cell lines, BxPC3, using chloroquine. Chloroquine is an endolytic agent that has been shown to enhance the endosomal escape of CPPs^62^ and RNA therapeutics^63,64^ by disrupting endosome integrity once it gets protonated at lower pH.^107–110^ Surprisingly, we did not observe lower CP_50_ values (which would signify improved endosomal escape) for any compounds in BxPC3 cells in the presence of chloroquine. In fact, there was a significant increase in CP_50_ for ct-SAHB and ct-nusinersen (**SI Figure 15**). There have been other instances where addition of chloroquine either barely enhanced or even reduced DNA delivery and gene expression.^111,112^ Chloroquine is also a lysosmotropic agent that can alter lysosomal pH and can prevent material from getting degraded. This raises the possibility that part of the CAPA signal derives from degraded material, and ct-SAHB and ct-nusinersen are more protected from degradation in the presence of chloroquine. While the CAPA data for these compounds do not resemble cases in which we have documented degradation artifacts,^45,58^ it remains a possibility that cannot be ruled out. Another explanation is that chloroquine has been shown to inhibit clathrin-mediated endocytosis,^33,113^ which our data (**Figure 5**, **SI Figure 14**) indicate is particularly important for cell penetration of polycationic peptides and nusinersen. Unlike chloroquine, the oligonucleotide escape enhancer compound UNC10217938A improved cell penetration in several cell lines tested, including primary HUVECs. Thus, such selective compounds may be more fruitful avenues for continued efforts in enhancing cell penetration of oligonucleotide therapeutics.

The experiments described in this work use standard cell culture protocols and will be easily expanded to additional cultured cell lines and primary cells of interest. The AAV plasmids are publicly available from Addgene, further facilitating the use of CAPA for cell-line-specific applications. We anticipate that these methods will be broadly useful for drug development, especially for peptide and oligonucleotide therapeutics targeting very specific tissues or cellular populations.

## Acknowledgments

This work was supported in part by NIH GM148407. The authors thank Dr. Mike Hanson, Director of the DNA/Peptide Facility at the University of Utah, for synthesis of nusinersen. BioRender was used for portions of the TOC graphic.

