## Supplemental Information for "Comparing cell penetration of biotherapeutics across human cell lines"

### Table of Contents

**Methods** 3

**SI Figure 1.** Chemical structures. 3

**SI Figure 2.** Structures of OECs. 7

**SI Figure 3.** Transduction of cell lines with AAV-H2GH. 7

**SI Figure 4.** HaloTag labeling with serial dilutions of ct-TAMRA in the seven cell lines. 8

**SI Figure 5.** CAPA dose-response curves for ct-compounds after 4h incubation (narrow gate). 9

**SI Figure 6.** CAPA dose-response curves for ct-compounds after 24h incubation (narrow gate). 9

**SI Figure 7.** CAPA dose-response curves for ct-compounds after 4h incubation (wide gate). 10

**SI Figure 8.** CAPA dose-response curves for ct-compounds after 24h incubation (wide gate). 10

**SI Figure 9**. $CP_{50}$ values for ct-compounds analyzed using the wide gate for seven cell lines. 11

**SI Figure 10.** Comparing CAPA data (4 hours) and $SAT_{50}$ of ct-TAMRA. 12

**SI Figure 11.** Comparing CAPA data (24 hours) and $SAT_{50}$ of ct-TAMRA. 13

**SI Figure 12.** Comparing $CP_{50}$ of ct-W and HaloTag expression. 14

**SI Figure 13.** Uptake of markers of endocytosis pathways at 37°C and 4°C in the 7 cell lines. 15

**SI Figure 14.** Comparison of nuclear penetration ($CP_{50}$) and levels of endocytosis pathways. 16

**SI Figure 15.** Effect of chloroquine in cell penetration in BxPC3 cells**.** 17

**SI Figure 16.** CAPA dose-response curves in BxPC3 cells with chloroquine. 17

**SI Figure 17.** CAPA dose-response curves in BxPC3 cells with or without OECs. 18

**SI Figure 18.** CAPA $CP_{50}$ values in BxPC3 and U2OS cells with or without OECs for ct-W. 18

**SI Figure 19.** CAPA dose-response curves in U2OS cells with or without OECs. 18

**SI Figure 20.** Clathrin knockdown validation in MIA-PaCa-2 cells. 19

**SI Figure 21***.* CAPA dose-response curves in MIA-PaCa2 cells after clathrin knockdown. 19

**SI Figure 22.** Uptake of markers of endocytosis pathways in the HUVEC primary cells. 20

**SI Figure 23.** Transduction of primary HUVEC cells. 20

**SI Figure 24.** CAPA dose-response curves in HUVEC cells with or without OECs. 20

**SI Table 1.** Culture media for each cell line. 4
**SI Table 2.** Transduction protocol for each cell line. 5
**SI Table 3.** Sequences and molecular weights of the panel of compounds used in this study. 21
**SI Table 4.** Statistical differences in $CP_{50}$ values for ct-R9W in the seven cell lines. 21
**SI Table 5.** Statistical differences in $CP_{50}$ values for ct-Tat in the seven cell lines. 22
**SI Table 6.** Statistical differences in $CP_{50}$ values for ct-SAHB in the seven cell lines. 23
**SI Table 7**. Statistical differences in $CP_{50}$ values for ct-W (4hours) in the seven cell lines. 24
**SI Table 8**. Statistical differences in $CP_{50}$ values for ct-nusinersen in the seven cell lines. 25
**SI Table 9**. Statistical differences in $CP_{50}$ values for ct-PMO in the seven cell lines. 26
**SI Table 10**. Statistical differences in $CP_{50}$ values for ct-R9W (24hours) in the seven cell lines. 27
**SI Table 11**. Statistical differences in $CP_{50}$ values for ct-W (24hours) in the seven cell lines. 28
**SI Table 12**. $CP_{50}$ values for ct-compounds in the seven cell lines (using the narrow gate). 29
**SI Table 13**. $CP_{50}$ values for ct-compounds in the seven cell lines (using the wide gate). 29
**SI Table 14.** Relative fluorescence values of endocytosis markers in the seven cell lines. 29
**SI Figure 15.** CAPA $CP_{50}$ values in BxPC3 cells with or without chloroquine. 30
**SI Table 16.** CAPA $CP_{50}$ values in BxPC3 cells with or without OECs. 30
**SI Table 17.** CAPA $CP_{50}$ values in U2OS cells with or without OECs. 30
**SI Table 18.** Statistics for ct-nusinersen (24h) $CP_{50}$ values in BxPC3 cells with or without OECs. 31
**SI Table 19.** Statistics for ct-PMO (24h) $CP_{50}$ values in the BxPC3 cells with or without OECs. 31
**SI Table 20.** Statistics for ct-nusinersen (24h) $CP_{50}$ values in the U2OS with or without OECs. 32
**SI Table 21.** Statistics for ct-PMO (24h) $CP_{50}$ values in the U2OS cells with or without OECs. 32
**SI Table 22.** Statistics in $CP_{50}$ for ct-W (24h) values in the BxPC3 cells with or without OECs. 33
**SI Table 23.** Statistics for ct-W (24h) in $CP_{50}$ values in the U2OS cells with or without OECs. 33
**SI Table 24**. CAPA $CP_{50}$ values in MIA PaCa-2 before or after clathrin knockdown. 34
**SI Table 25.** Unpaired t-test comparing the markers uptake before and after clathrin knockdown in MIA-PaCa-2 cells. 34
**SI Table 26.** Unpaired t-test comparing the $CP_{50}$s of ct-compounds before and after clathrin knockdown in MIA-PaCa-2 cells. 34
**SI Table 27.** CAPA $CP_{50}$ values in HUVEC cells with or without OECs. 34

### Methods

#### Synthesis of chloroalkane tagged (ct-) amine (ct-$NH_2$), dibenzylcyclooctyne (ct-DBCO), and tetramethylrhodamine (ct-TAMRA)

The structures of ct-$NH_2$, ct-DBCO, and ct-TAMRA are shown in **SI Figure 1**. They were synthesized as previously described[1] and their identity and purity were confirmed by MALDI-TOF mass spectrometry and reverse phase high performance liquid chromatography (RP-HPLC), respectively.

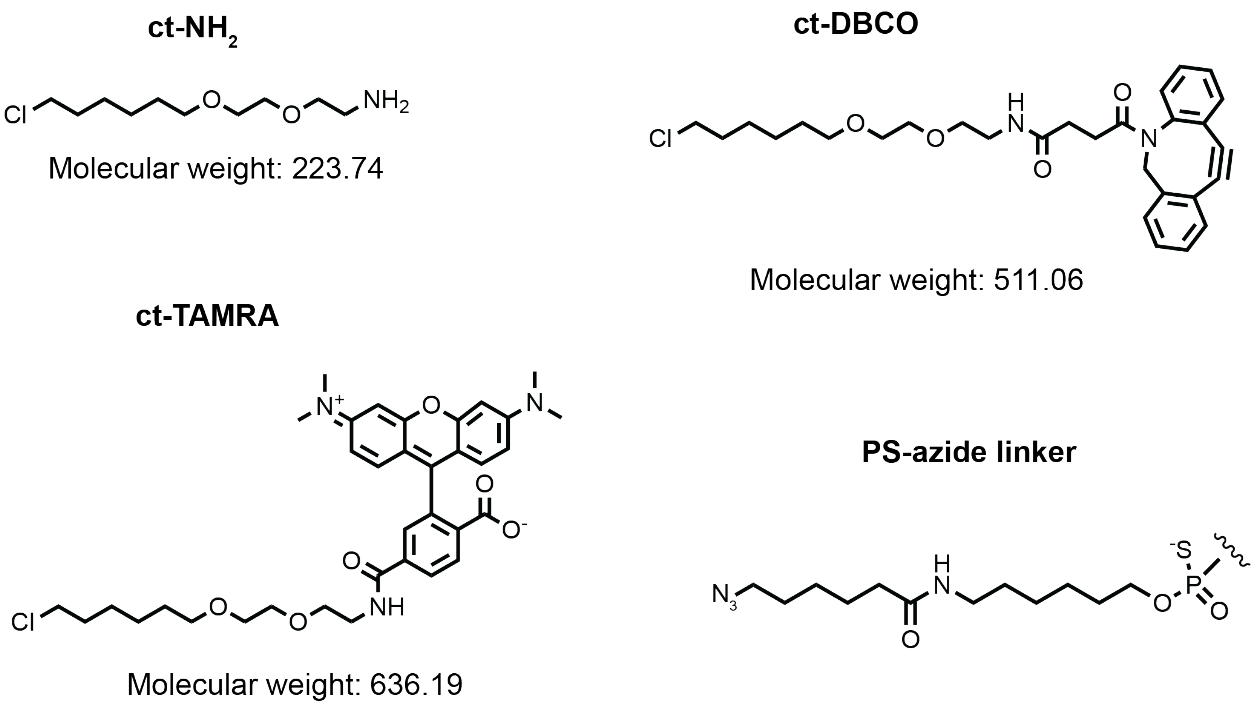


**SI Figure 1. Chemical structures.** Structures and molecular weights of ct-$NH_2$, ct-DBCO, and ct-TAMRA, and structure of PS-azide linker on nusinersen.

#### Peptide synthesis and purification

Peptides were synthesized on Rink Amide Resin by standard Fmoc chemistry using the Prelude peptide synthesizer by Gyros Protein Technologies. For each coupling, 5 equivalents of Hydroxybenzotriazole (HOBt), 10 equivalents of 1H-Benzotriazole-1-yl)-1,1,3,3-tetramethyluronium hexafluorophosphate (HBTU), and 10 equivalents of diisopropylethylamine (DIPEA) were dissolved in *N*,*N*-dimethylformamide (DMF) and shaken with the resin for 20 min. Deprotection used 20% piperidine in DMF. To add the chloroalkane tag to the N-terminus of each peptide, 3 equivalents of chloroalkane carboxylic acid (RL-3180, Iris Biotech), 3 equivalents of benzotriazol-1-yl-oxytripyrrolidinophosphonium hexafluorophosphate (PyBOP), and 10 equivalents of DIPEA were dissolved in DMF, added to the resin (after Fmoc deprotection) and shaken at room temperature for 1 h. For the stapled peptide, SAHB, an olefin metathesis reaction was performed according to the literature to staple the two *(S)*-2-(4′-pentenyl)alanines.[2] Briefly, 8 mg of Grubbs catalyst, generation 1 were dissolved in 2 mL 1,2-dichloroethane (DCE), added to resin and shaken for 3 h. After the reaction solution was removed and the resin was washed, fresh Grubbs catalyst solution was added and the reaction was shaken overnight to maximize yield. The peptides were cleaved from the resin using a cleavage cocktail (94% trifluoroacetic acid (TFA), 2.5% ethanedithiol (EDT), 2.5% $H_2O$, and 1% trrsopropylsilane (TIPS)) and precipitated in ice-cold diethyl ether. Peptides were dissolved in 50:50 water: acetonitrile solution and purified by RP-HPLC and lyophilized. Peptide stocks in dimethyl sulfoxide (DMSO) were made and the concentration was measured using their UV absorbance at 280 nm.

**Conjugation of ct-DBCO to oligonucleotides and purification of ct-oligonucleotides**
Nusinersen was synthesized by the DNA/Peptide Facility, part of the Health Sciences Center Cores at the University of Utah, and the PMO was synthesized by Gene Tools. Both oligonucleotides were synthesized with a 5′ azide group. Lyophilized oligos were dissolved in 100 mM aqueous triethylamine acetic acid (TEAAc) pH 7 with 30% acetonitrile (nusinersen) or water (PMO). Two equivalents of ct-DBCO were added and the reaction was stirred overnight at room temperature. The mixture was purified by RP-HPLC. The mobile phase was a gradient of 95% solvent A (5% acetonitrile in 100 mM aqueous TEAAc, pH 7) and 5% solvent B (20% 100 mM aqueous TEAAc, pH 7 and 80% acetonitrile) to 100% solvent B in 20 minutes. The identity of the ct-oligonucleotides was confirmed by MALDI-TOF mass spectrometry, and the purity was verified by analytical RP-HPLC using an XBridge Oligonucleotide BEH C18 column and the gradient described above. The pure fractions were lyophilized, desalted, and stored at -20°C in PBS (ct-nusinersen) or at room temperature in water (ct-PMO, according to GeneTool's recommendations).

**Cell culture and maintenance**
Cell lines were acquired from ATCC and their identities were verified using ATCC's authentication services (Human Cell STR profiling, 135-XV). Every cell line was tested for mycoplasma using a mycoplasma testing kit (ATCC 3012K). Cell lines were cultured according to ATCC's recommendations (see **SI Table 1**). All media were acquired from ATCC and fetal bovine serum (FBS) was acquired from Neuromics.

**SI Table 1. Culture media for each cell line.**

| **Cell line** | **Culture media** |
|---|---|
| HeLa | DMEM + 10% FBS |
| Saos2 | McCoy's 5a medium modified + 15% FBS |
| U2OS | McCoy's 5a medium modified + 10% FBS |
| MIA-PaCa2 | DMEM + 10% FBS + 2.5% horse serum |
| HEK293T | DMEM + 10% FBS (heat inactivated) + 2mM L-glutamine |
| BxPC3 | RPMI-1640 + 10% FBS |
| HepG2 | EMEM + 10% FBS |
| HUVEC | Vascular Cell Basal medium supplemented with endothelial cell growth kit- VEGF |

**Transduction of cell lines**
Cells were seeded at a density of 1 x $10^4$ cells per well in 96-well tissue culture treated plates and 16-24 h after, they were transduced with AAV2 particles expressing a histone 2B-GFP-HaloTag fusion.[3] The vector, available at Addgene (#182202) was packaged in AAV using the services offered by the Salk Institute or by Vector Builder. The transduction protocol followed for each cell line is found in **SI Table 2**. AAV2 particles were diluted in the respective media at a specific multiplicity of infection (MOI) and 125 μL of the mixture was added per well. The cells were incubated at 37°C, 5% $CO_2$ with the AAV for 24 or 48 h. Transduction protocols were optimized for each cell line for robust and similar HaloTag expression.

**SI Table 2. Transduction protocol for each cell line.**

| Cell line | Transduction protocol | |
|---|---|---|
| | MOI | Transduction time |
| HeLa | $10^4$ | 24 h |
| Saos2 | $10^4$ | 24 h |
| U2OS | $10^3$ | 24 h |
| MIA-PaCa2 | $5 \times 10^4$ | 48 h |
| HEK293T | $10^4$ | 24 h |
| BxPC3 | $3 \times 10^5$ | 48 h |
| HepG2 | $10^6$ | 48 h |
| HUVEC | $10^6$ | 48 h |

**Chloroalkane Penetration Assay (CAPA)**

CAPA was performed as described previously.[1,3,4] Briefly, for each cell line, compounds were diluted in optiMEM and incubated with cells at 37°C, 5% $CO_2$ for 4 or 24 h (pulse step). After the pulse step, media was aspirated and cells were washed with 80 μL optiMEM for 15 min at room temperature. Upon aspiration of the optiMEM, cells were treated with 50 μL of 5 μM ct-TAMRA for 15 min at room temperature (chase step), and then washed briefly with 80 μL of optiMEM at room temperature. Cells were then trypsinized with 40 μL of 0.05% trypsin in PBS, suspended in an additional 180 μL PBS and analyzed by flow cytometry.

For each experiment, three wells were treated with only optiMEM during the pulse and chase steps, serving as the 0% fluorescence control. Three additional wells were treated with only optiMEM during the pulse step and ct-TAMRA during the chase step, serving as the 100% fluorescence control. Also, a row of cells was pulsed only with optiMEM, then treated with serial dilutions of ct-TAMRA during the chase step in order to generate a HaloTag saturation curve for each cell line. Three biological trials were performed for each CAPA experiment. Data points in figures reporting CAPA data are shown as averages with error bars that represent the standard error of the mean from the three independent trials. All $CP_{50}$ values are reported as averages and standard errors from three independently calculated curve fits from the three independent trials.

**Endocytosis assays using fluorescently labeled markers**

Cells were seeded at a density of $1 \times 10^4$ cells per well in a 96-well plate and one day later, they were treated with fluorescently labeled markers. Specifically, they were treated with 50 μL of 1 mg/mL TMR-dextran (ThermoFisher Scientific D1819), 1 mg/mL TMR-transferrin (ThermoFisher Scientific T2872), or 5 μg/mL Alexa Fluor™ 647 cholera toxin subunit B (ThermoFisher Scientific C34778); 50 μL optiMEM was used as a control. The cells were incubated with the markers for 1 h at either 37°C, 5% $CO_2$ or at 4°C. After aspiration, the cells were washed with 80 μL of optiMEM for 30 min at room temperature. Cells were trypsinized with 40 μL of 0.05% trypsin in PBS, suspended in an additional 180 μL PBS and analyzed by flow cytometry. Values for relative fluorescence units (RFU) were calculated by dividing the mean fluorescence units of each sample by the mean fluorescence units of the background. At least three biological trials were performed for each cell line and RFU values are shown as averages with standard error of the mean from the three independent trials.

**Clathrin knockdown**

MIA-PaCa2 cells were seeded at a density of 1.5 x $10^5$ cells per well in a 6-well plate (day 1). On day 2, cells were transduced with 2 mL of AAV at MOI of $10^4$. On day 3, media was aspirated, and cells were transfected with 10 nM siRNA targeting *clathrin heavy chain* using transfection reagent DharmaFECT 1 from Dharmacon. siRNA were designed using ON-TARGET plus SMART pool for Human CLTC (L-004001-01) and ON-TARGET plus non-targeting pool (D-001810-10-05) and were obtained from Dharmacon. The transfection was carried out as follows: 200 μL of 10 nM siRNA (clathrin-targeting or scrambled) in optiMEM were added in Tube 1 and 10 μL DharmaFECT 1 in 190 μL optiMEM were added in Tube 2. After incubating for 5 min at room temperature, contents of Tube 1 were added to Tube 2 and the mixture was incubated for 20 min at room temperature. After that, 1,600 μL of full media with FBS was added. Old media was aspirated from cells and the 2,000 μL were added to the cells. After 72 hours of incubation, cells were washed with 1 mL PBS and then trypsinized with 500 μL 0.05% trypsin/PBS and suspended in an additional 1 mL PBS. Cells were spun down at 1,000rpm for 3 minutes and after removing the supernatant they were either lysed and used for Western blots or re-seeded in 96-well plates for CAPA and endocytosis marker experiments.

**Western blots**

Cells were lysed in 60 μL RIPA buffer by undergoing ten freeze-thaw cycles in liquid nitrogen and a room temperature water bath. Lysates were centrifuged at 4°C for 20 minutes at >13,500rpm and the supernatant was transferred to clean tubes. The amount of protein was quantified using a BSA standard curve and the Pierce™ 660 nm protein assay reagent from ThermoFisher Scientific. From each lysate, 15 μg of total protein was separated on a 4-15% SDS-PAGE gel and transferred to a nitrocellulose membrane using the iBlot 2 PVDF mini stacks and the iBlot2 system (ThermoFisher). The membrane was blocked in StartingBlock™ (PBS) blocking buffer (ThermoFisher) for 30 min at room temperature and then transferred to an iBind™ flex card (Invitrogen) with 1:200 anti-clathrin rabbit IgG (PA5-25804, ThermoFisher) or with 1:1000 anti-GAPDH rabbit IgG (D16H11, Cell Signaling Technology). Membranes were then exposed to 1:1000 anti-rabbit IgG fused to HRP (A16110, ThermoFisher). The membrane was incubated with the antibodies usind the iBind system (ThermoFisher) for 4 hours and then washed in TBST for 5 minutes and in TBS 3x for 3mins each, and then incubated with SuperSignal™ West Pico chemiluminescent substrate (ThermoFisher) and developed using a ChemiDoc MP imaging system (BioRad).

**Small molecule enhancers**

The three small molecule enhancers of endosomal escape are shown in **SI Figure 2**. 60 μM chloroquine, 10 μM UNC10217938A, or 5 μM SH-BC-893 were used according to existing literaute.[5–7] The compounds were co-incubated with the ct-compounds for the desired incubation time (4h or 24 h).

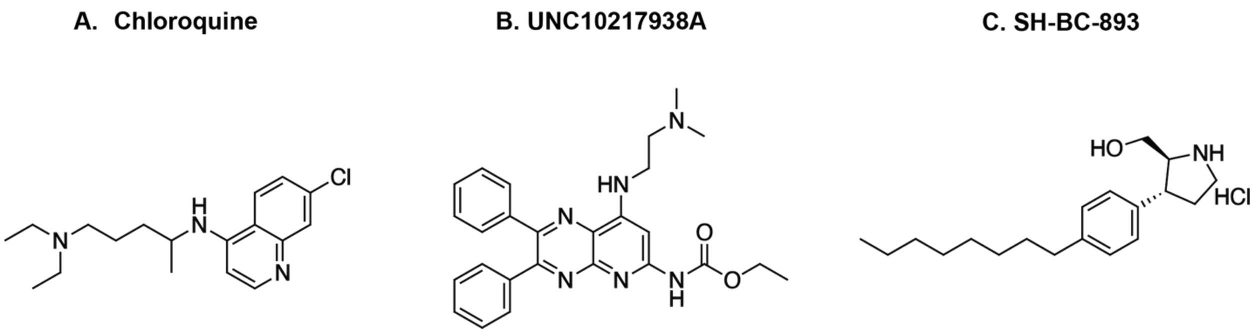


**SI Figure 2**. **Structures of OECs.** Small molecule enhancers of endosomal escape.

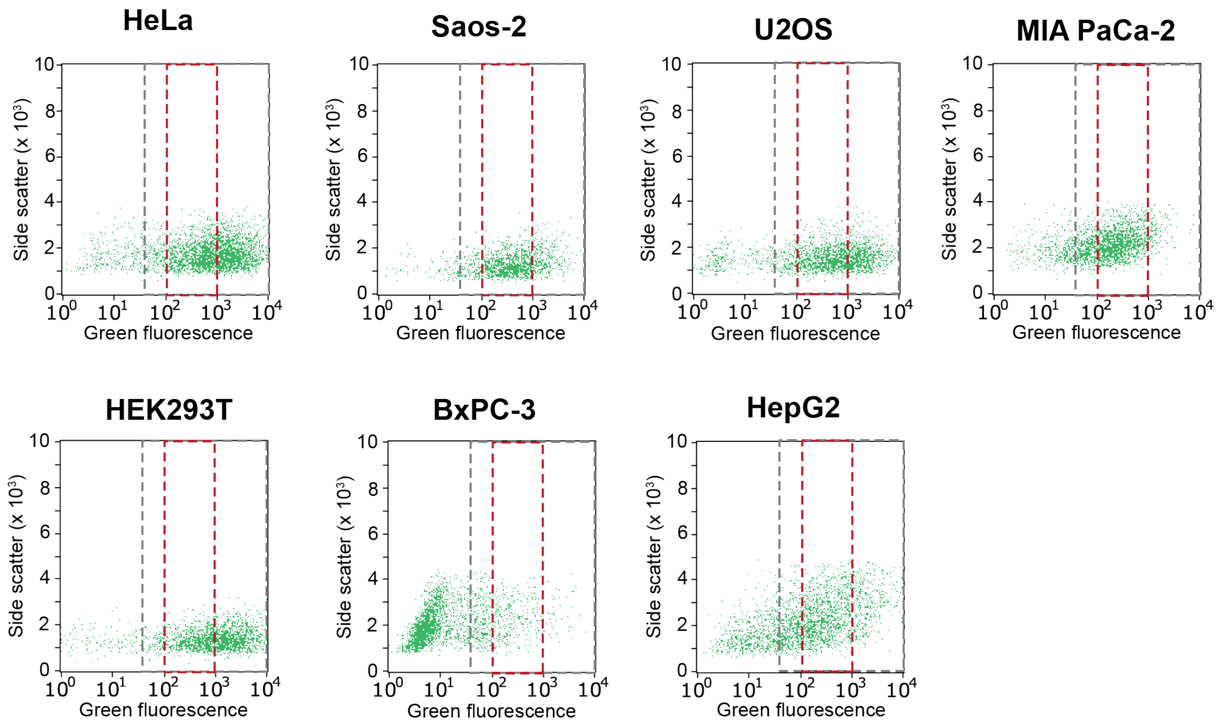


**SI Figure 3. Transduction of cell lines with AAV-H2GH.** Flow cytometry plots showing side scatter versus green fluorescence for the seven cell lines. Grey rectangles indicate “wide” gates and red rectangles indicate “narrow” gates calibrated for positive green fluorescence to analyze only cells expressing HaloTag-GFP.

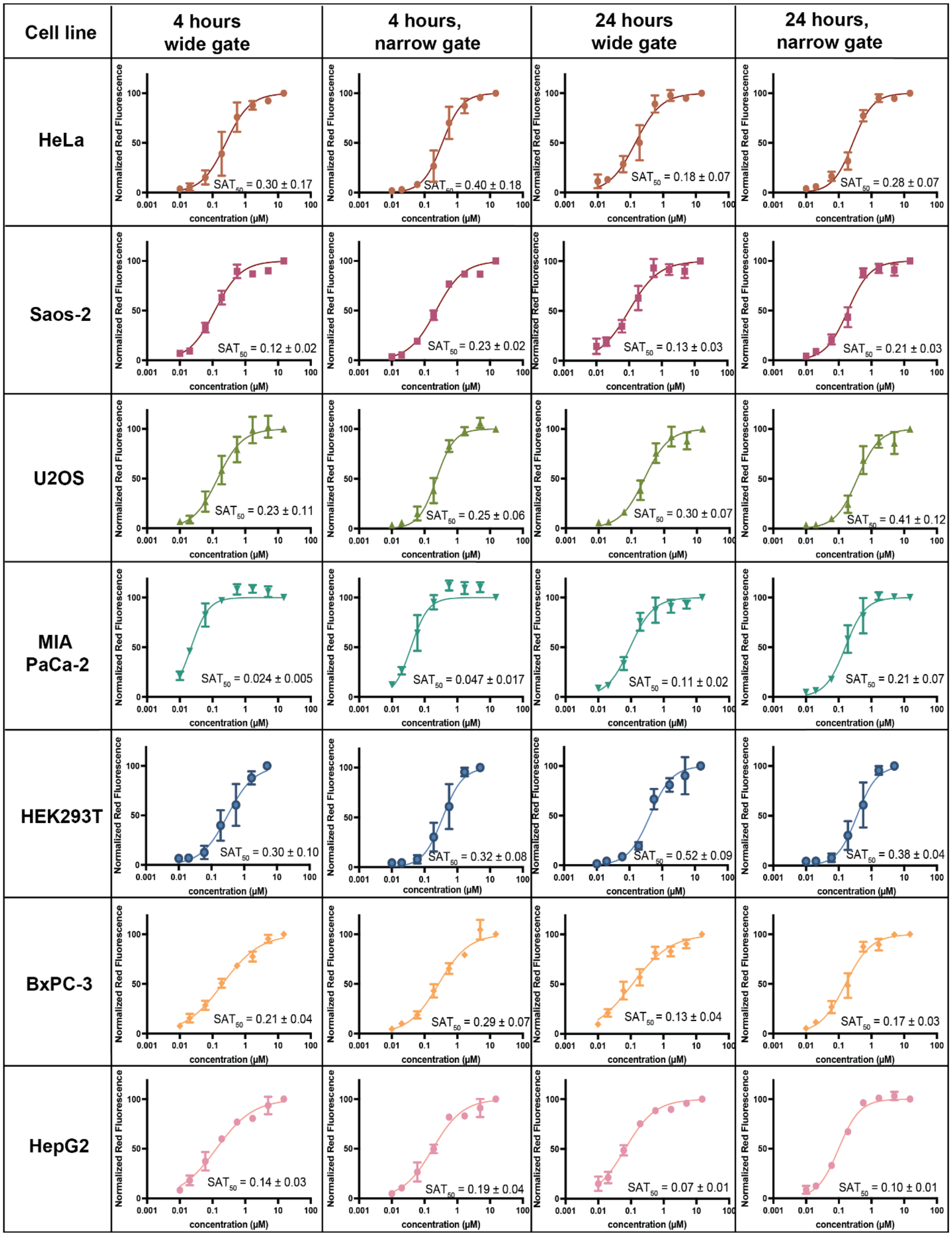


**SI Figure 4. HaloTag labeling with serial dilutions of ct-TAMRA in the seven cell lines.** CAPA experiments were performed using 4h and 24h chase steps using two different gates, a “wide” gate and a “narrow” gate (**SI Figure 3**). At the higher concentrations of ct-TAMRA, HaloTag gets saturated with ct-TAMRA during the chase step. Curve fits are to a general $IC_{50}$ curve, the midpoint of which was defined as the $SAT_{50}$ for that cell line. Plots show results from three independent trials and error bars show standard error of the mean. $SAT_{50}$ values are reported as the average and standard error of the mean from three independent curve fits to three independent trials.

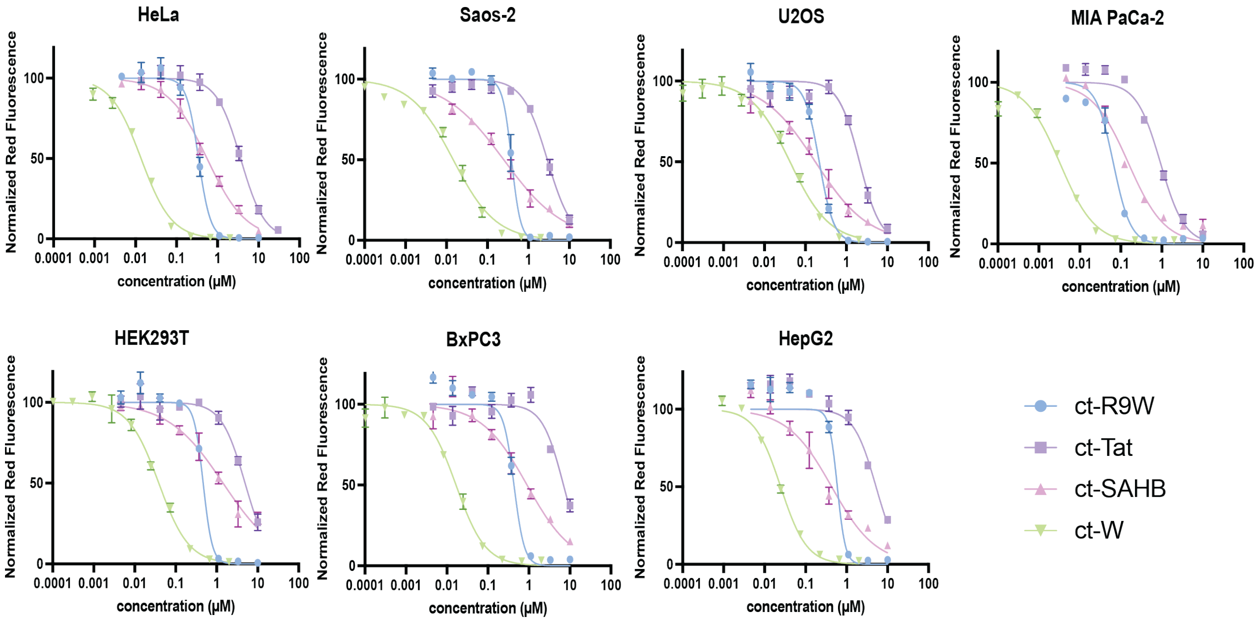


**SI Figure 5. CAPA dose-response curves for ct-compounds after 4 hours incubation in seven cell lines (narrow gate).** All data show results from three independent trials and error bars show standard error of the mean. $CP_{50}$s shown in **Figure 2A** and **SI Table 12**.

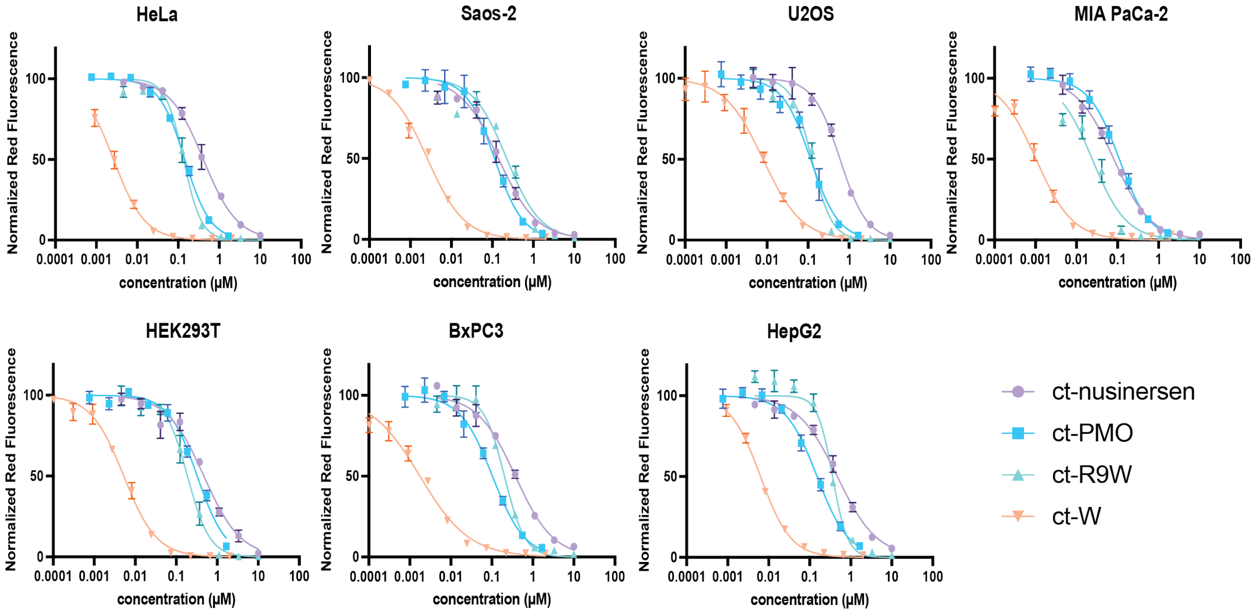


**SI Figure 6. CAPA dose-response curves for ct-compounds after 24 hours incubation in seven cell lines (narrow gate).** All data show results from three independent trials and error bars show standard error of the mean. $CP_{50}$s shown in **Figure 2B** and **SI Table 12.**

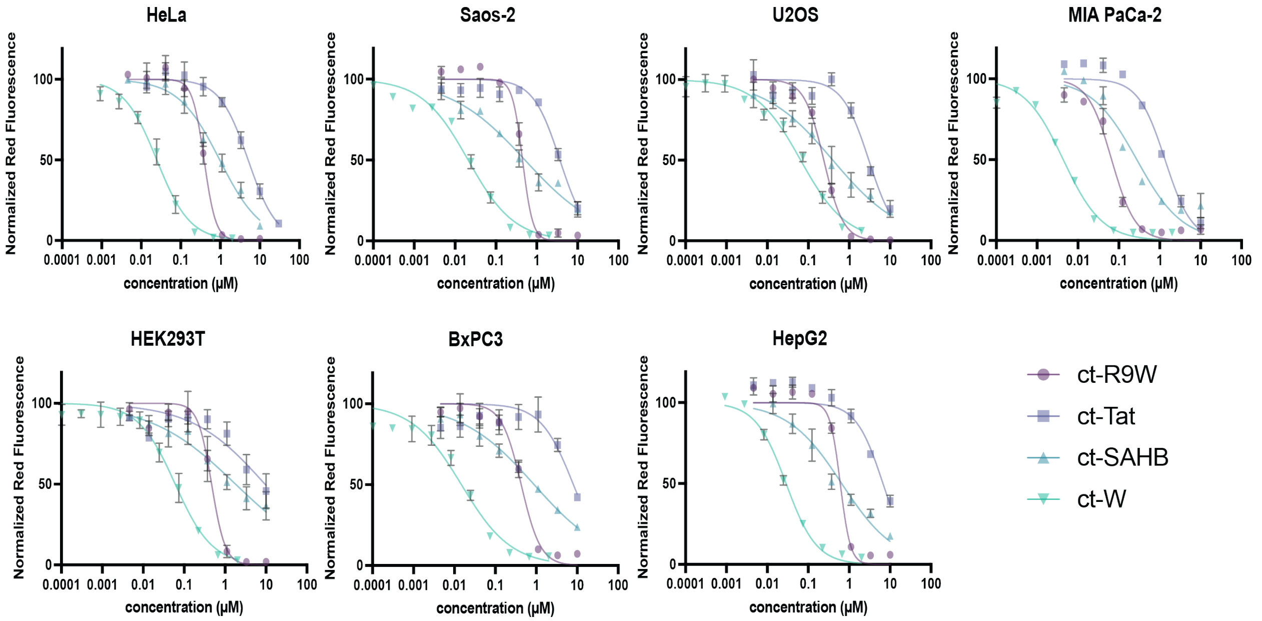


**SI Figure 7. CAPA dose-response curves for ct-compounds after 4 hours incubation in seven cell lines (<u>wide gate</u>).** All data show results from three independent trials and error bars show standard error of the mean. $CP_{50}$s shown in **SI Figure 9** and **SI Table 13**.

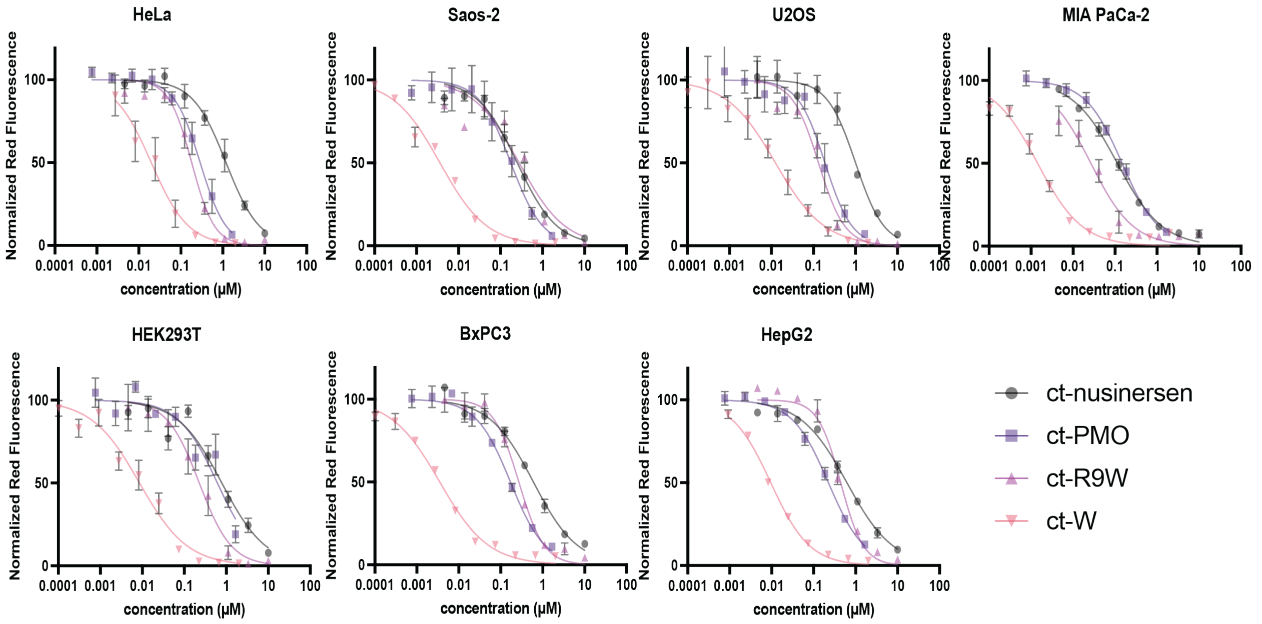


**SI Figure 8. CAPA dose-response curves for ct-compounds after 24 hours incubation in seven cell lines (<u>wide gate</u>).** All data show results from three independent trials and error bars show standard error of the mean. $CP_{50}$s shown in **SI Figure 9** and **SI Table 13**.

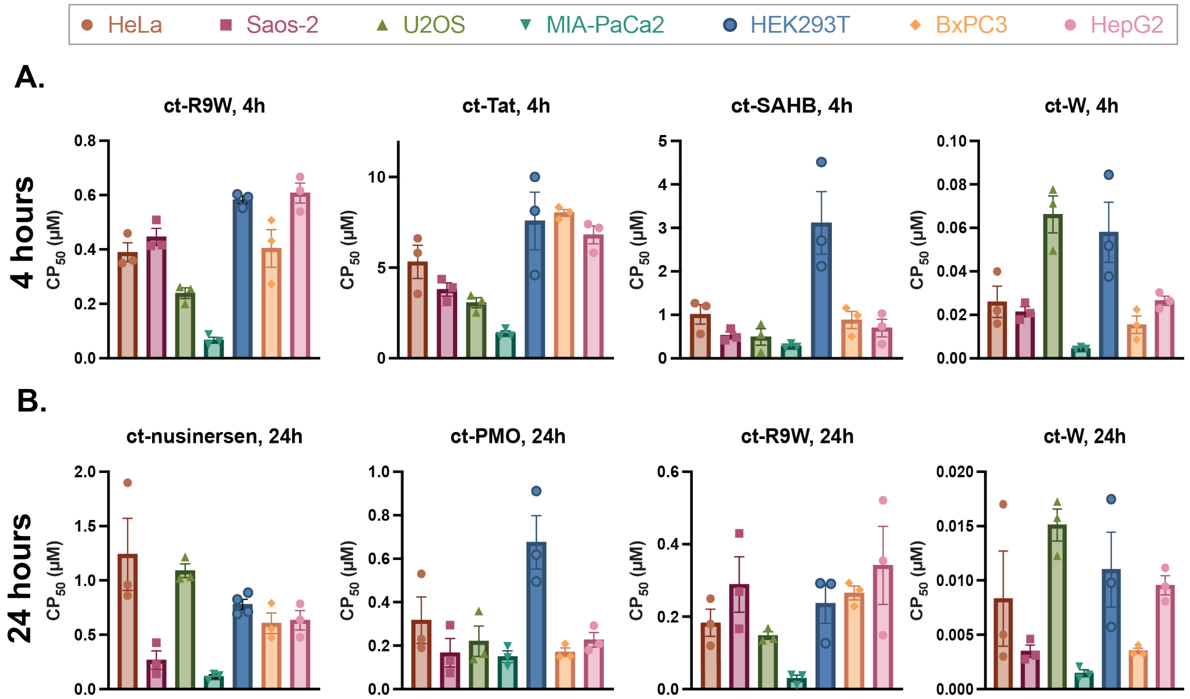


**SI Figure 9. CP$_{50}$ values for ct-compounds analyzed using the *wide gate* for seven cell lines. (A)** CP$_{50}$ values for each ct-compound in each cell line after 4 h incubation. These values are derived from curve fits shown in **SI Figure 7**. **(B)** CP$_{50}$ values for each ct-compound in each cell line after 24 h incubation. These values are derived from curve fits shown in **SI Figure 8**. CP$_{50}$ values are reported as the average and standard error of the mean from three independent curve fits to three independent trials.

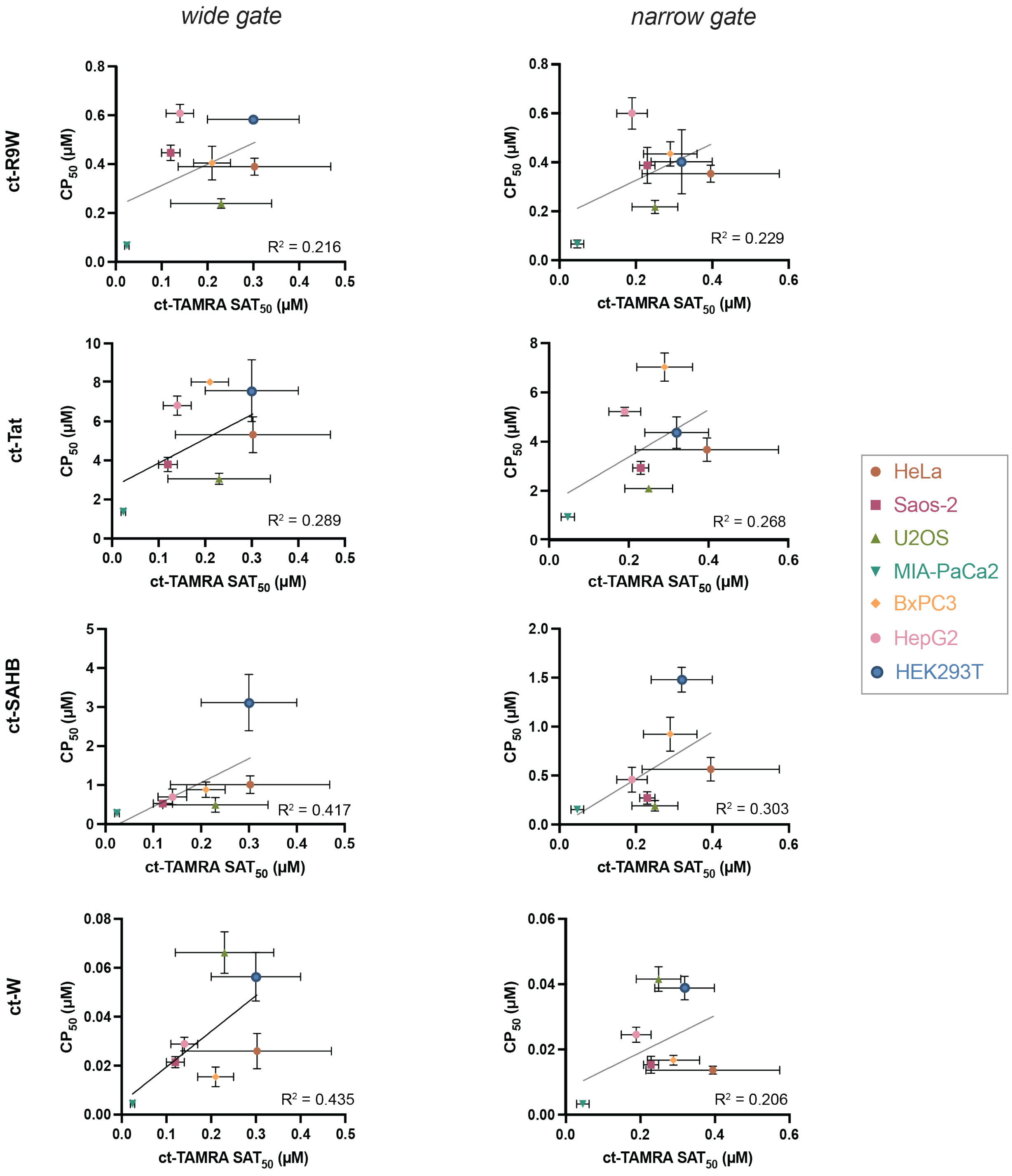


**SI Figure 10. Comparing CAPA data (4 hours) and $SAT_{50}$ of ct-TAMRA.** $CP_{50}$ and $SAT_{50}$ values are reported as the average and standard error of the mean from three independent curve fits to three independent trials.

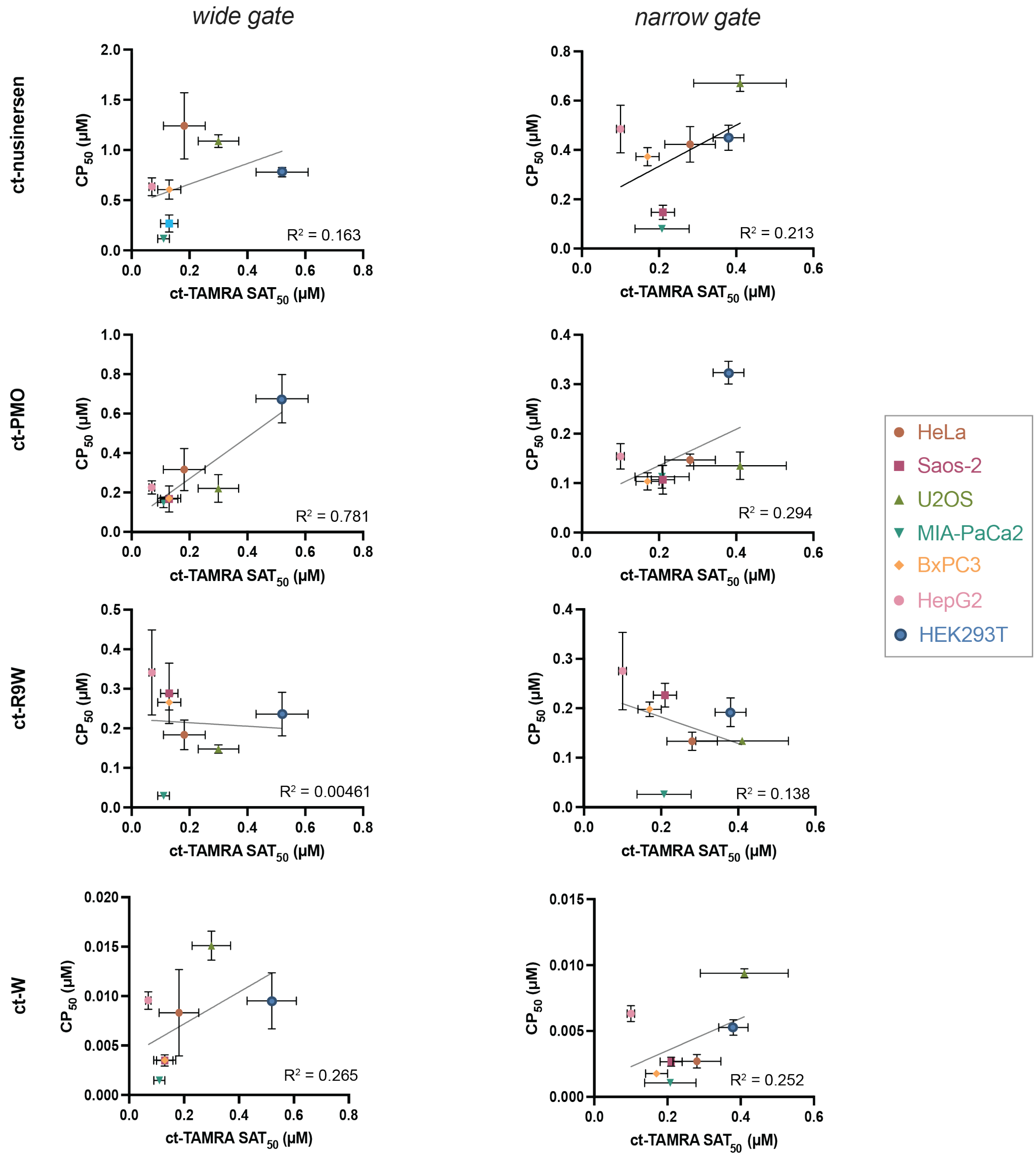


**SI Figure 11. Comparing CAPA data (24 hours) and $SAT_{50}$ of ct-TAMRA.** $CP_{50}$ and $SAT_{50}$ values are reported as the average and standard error of the mean from three independent curve fits to three independent trials.

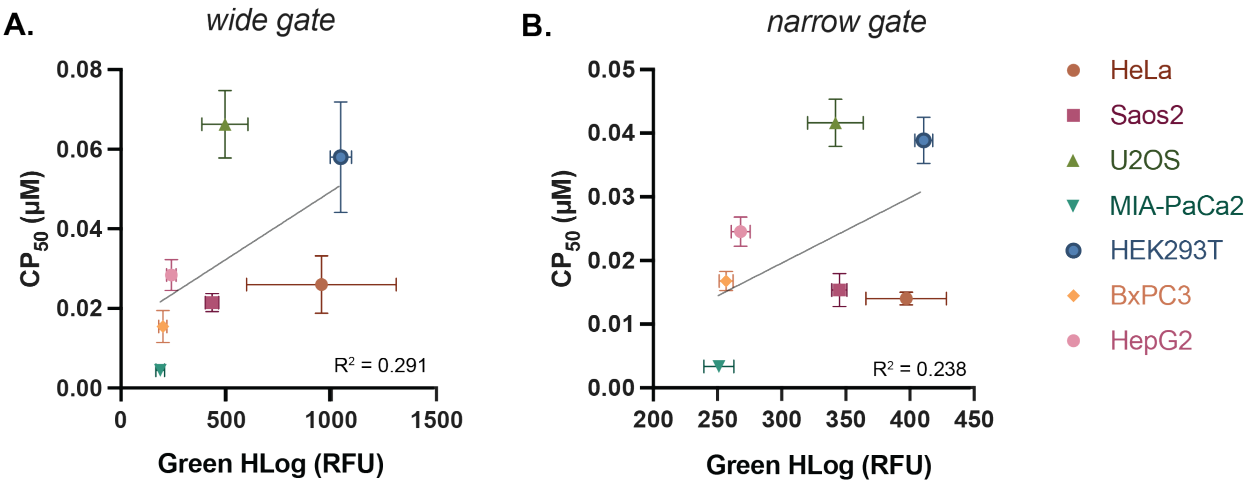


**SI Figure 12. Comparing $CP_{50}$ of ct-W and HaloTag expression.** Analysis using **(A)** a wide gate, **(B)** a narrow gate. $CP_{50}$ values are reported as the average and standard error of the mean from three independent curve fits to three independent trials. Green HLog values are reported as the average and standard error of the mean of the green fluorescence from cells that had not been treated with ct-compound or ct-dye in three independent trials.

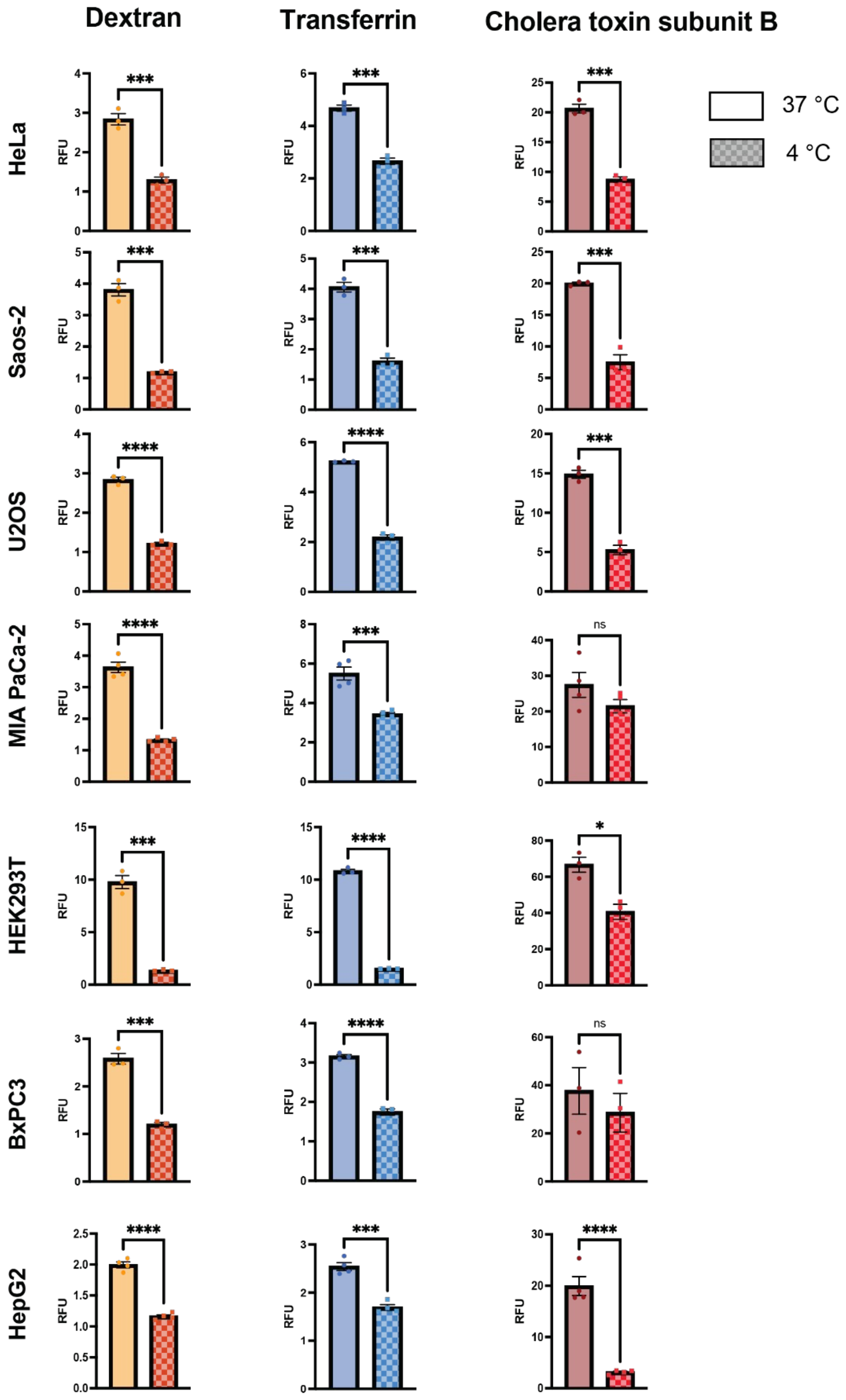


**SI Figure 13. Uptake of markers of endocytosis pathways at 37°C versus 4°C in the seven cell lines**. RFU values are reported as the average and standard error of the mean from three independent trials.

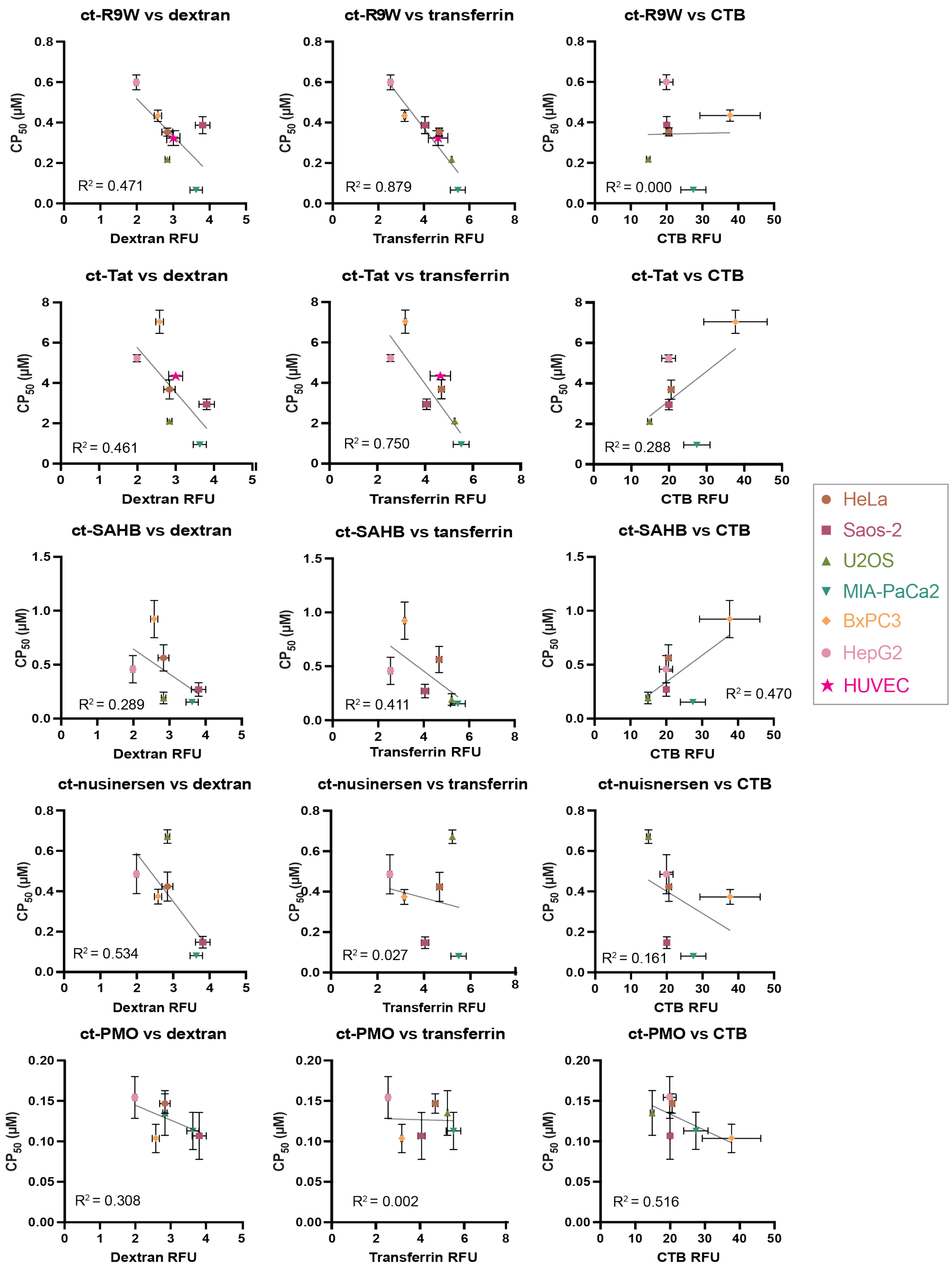


**SI Figure 14. Comparison of nuclear penetration ($CP_{50}$) and levels of endocytosis pathways.** Analysis for ct-R9W, ct-Tat, ct-SAHB, ct-nusinersen, and ct-PMO in six different cell lines. Note that the HUVEC data for ct-R9W and ct-Tat were not taken into account in the line equation. RFU and $CP_{50}$ values are reported as the average and standard error of the mean from three independent trials.

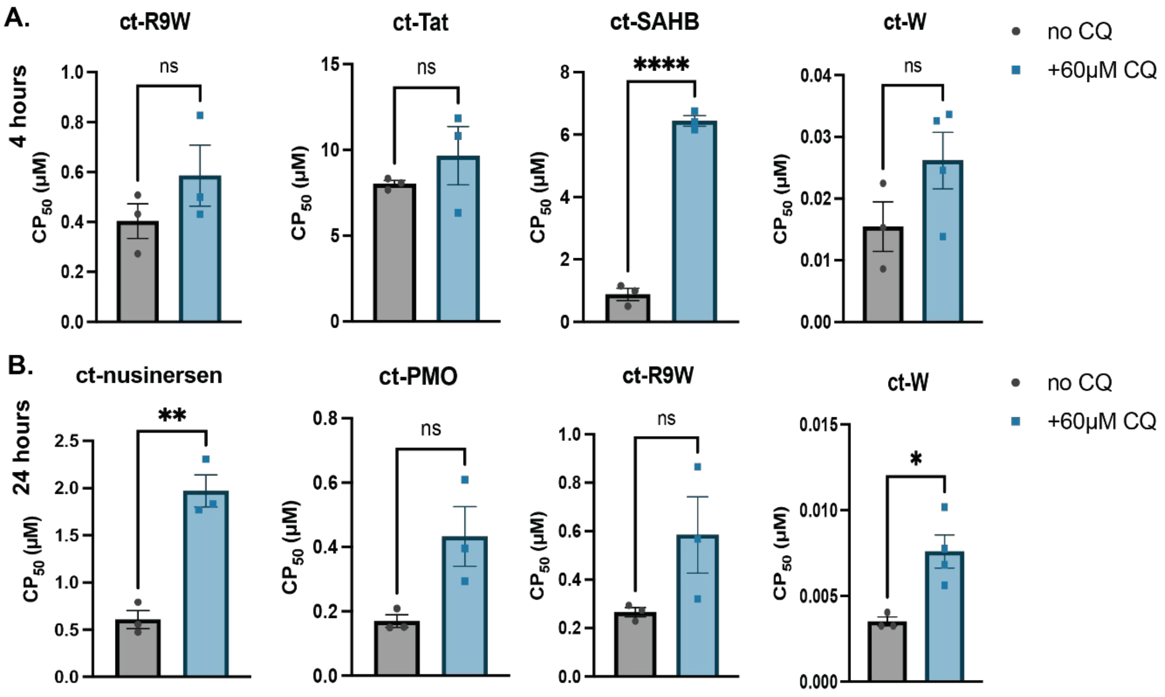


**SI Figure 15. Effect of chloroquine in cell penetration in BxPC3 cells**. $CP_{50}$ values for ct-compounds in the absence or presence of 60 μM chloroquine in **BxPC3 cells** after **(A)** 4 hours, or **(B)** 24 hours of co-incubation. These values are derived from curve fits shown in **SI Figure 16**. $CP_{50}$ values are reported as the average and standard error of the mean from three independent curve fits to three independent trials. An unpaired t-test was used to calculate *p* values $*p \leq 0.05$, $**p<0.01$, $****p \leq 0.001$.

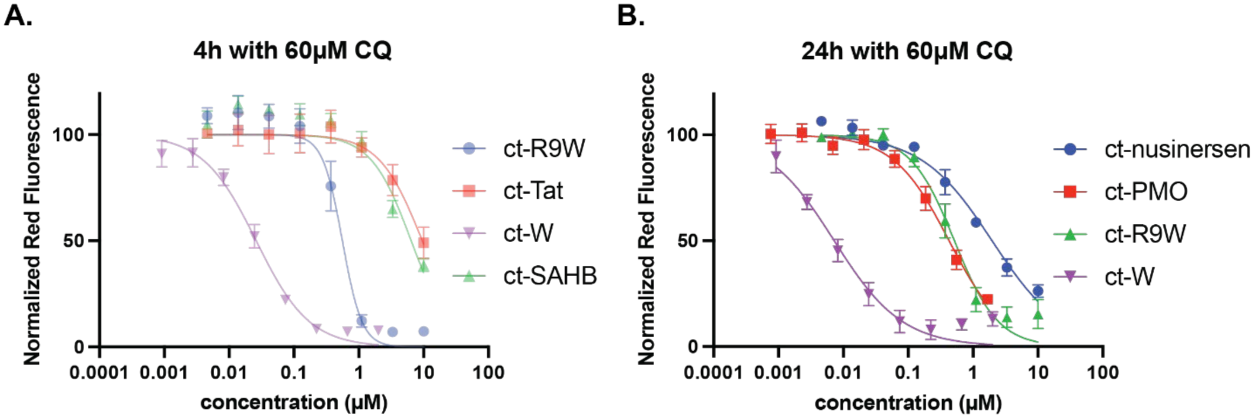


**SI Figure 16. CAPA dose-response curves in BxPC3 cells with chloroquine.** Co-incubation of ct-compounds with 60 μM chloroquine after **(A)** 4 hours, or **(B)** 24 hours in BxPC3 cells. All data show results from three independent trials and error bars show standard error of the mean. $CP_{50}$s shown in **SI Figure 15** and **SI Table 15**.

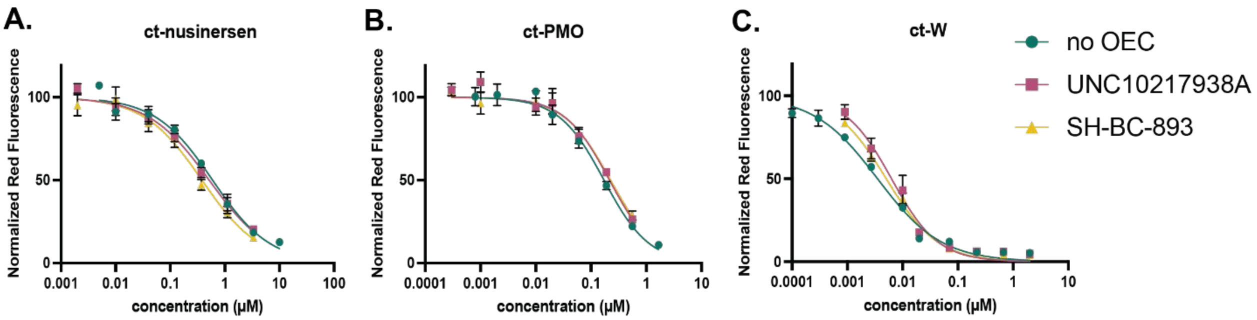


**SI Figure 17: CAPA dose-response curves in BxPC3 cells with or without OECs.** Data for **(A)** ct-nusinersen, **(B)** ct-PMO, **(C)** ct-W following 24 hours of incubation BxPC3 cells in the presence of 10 μM UNC10217938A, 5 μM SH-BC-893 or absence of oligonucleotide enhancer compound (no OEC). All data show results from three independent trials and error bars show standard error of the mean. $CP_{50}$ values shown in **Figure 4A**, **SI Figure 18A** and **SI Table 16**.

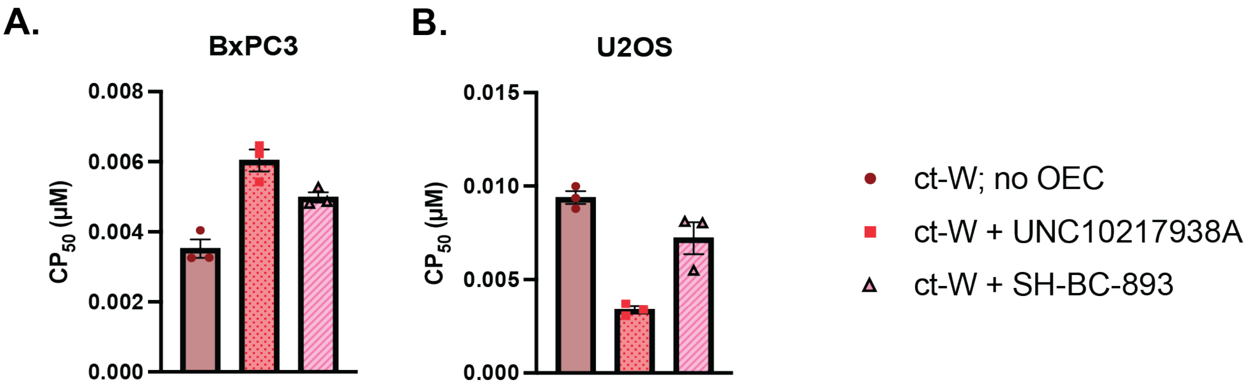


**SI Figure 18: CAPA $CP_{50}$ values in BxPC3 and U2OS cells with or without OECs for ct-W.** Co-incubation of 10 μM UNC10217938A and 5 μM SH-BC-893 and ct-W **(A)** in BxPC3 cells and **(B)** in U2OS cells for 24 hours. These values are derived from curve fits shown in **SI Figure 17, 19**. $CP_{50}$ values are reported as the average and standard error of the mean from three independent curve fits to three independent trials. Statistical differences for these data are found in **SI Tables 22,23**.

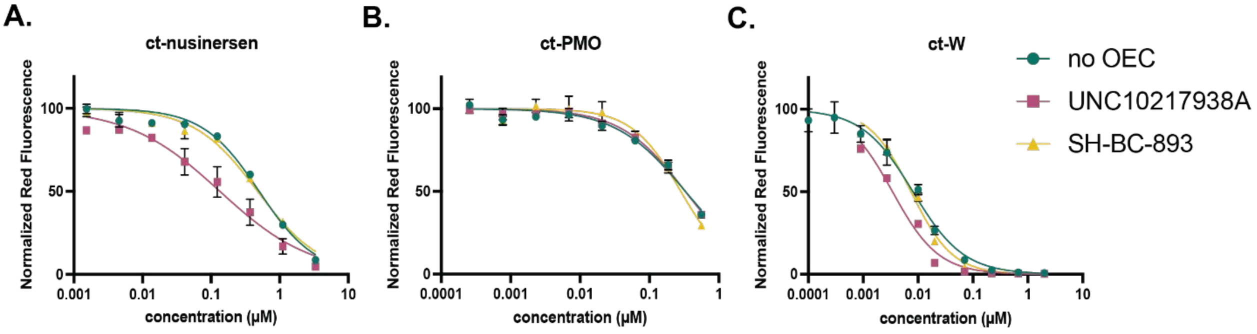


**SI Figure 19: CAPA dose-response curves in U2OS cells with or without OECs.** Data for **(A)** ct-nusinersen, **(B)** ct-PMO, **(C)** ct-W following 24 hours of incubation U2OS cells in the presence of 10μM UNC10217938A, 5μM SH-BC-893 or absence of oligonucleotide enhancer compound (OEC). All data show results from three independent trials and error bars show standard error of the mean. $CP_{50}$s shown in **Figure 4B**, **SI Figure 18B** and **SI Table 17**.

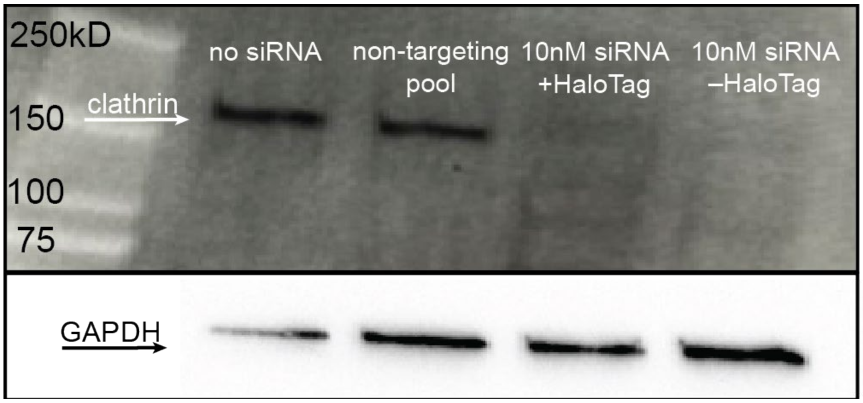


**SI Figure 20. Clathrin knockdown validation in MIA-PaCa-2 cells.** Anti-clathrin and anti-GAPDH Western blots for MIA PaCa-2 cells (no siRNA), with 10nM non-targeting pool siRNA, and 10 nM siRNA targeting clathrin heavy chain in cells that have been transduced with AAV (+HaloTag) or not (–HaloTag).

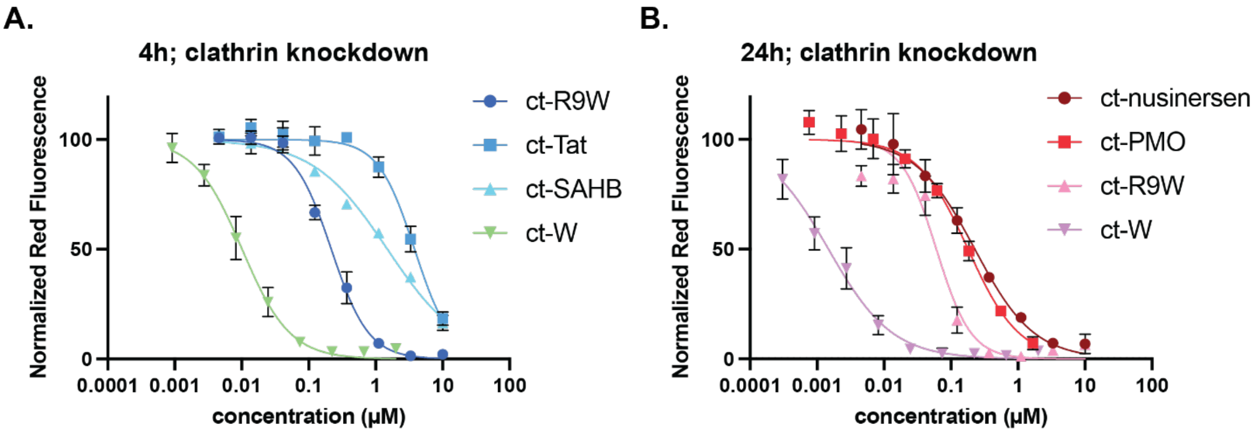


**SI Figure 21. CAPA dose-response curves in MIA-PaCa2 cells after clathrin knockdown.** Data after **(A)** 4 hours, or **(B)** 24 hours of incubation. All data show results from three independent trials and error bars show standard error of the mean. $CP_{50}$s shown in **Figure 5C & 5D and SI Table 24**.

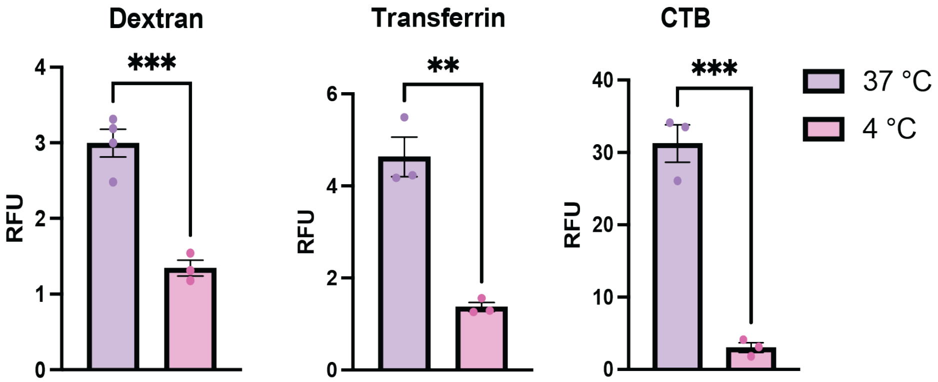


**SI Figure 22: Uptake of markers of endocytosis pathways in the HUVEC primary cells.** RFU values are reported as the average and standard error of the mean from three independent trials.

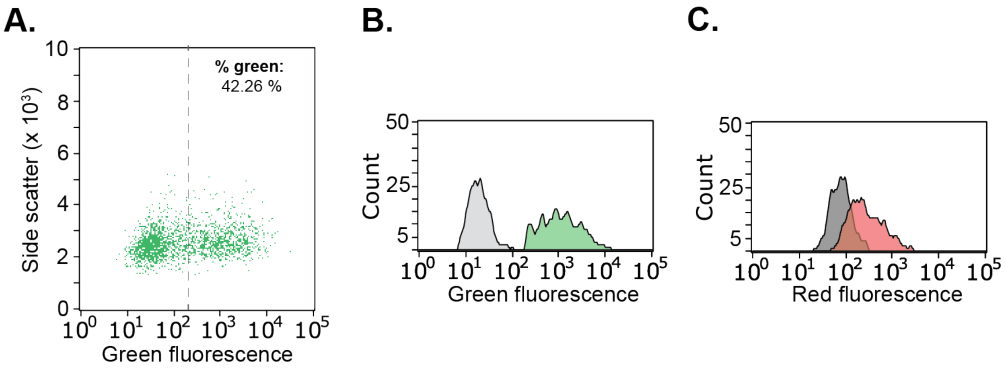


**SI Figure 23. Transduction of primary HUVEC cells**. **(A)** Flow cytometry plot showing side scatter versus green fluorescence for HUVEC cells. **(B)** Green fluorescence shift of transduced cells (green) versus non-transduced cells (light grey). **(C)** Shift in red fluorescence of transduced cells upon treatment with ct-TAMRA (red) or treatment with optiMEM (dark grey).

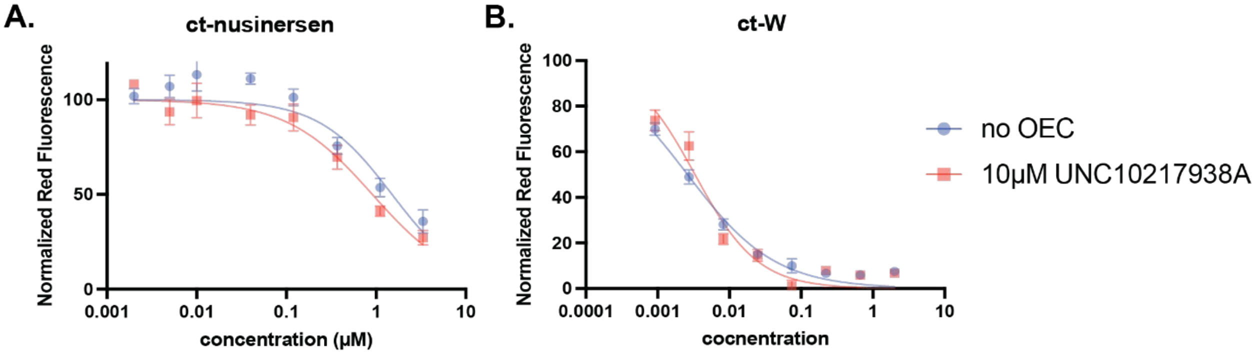


**SI Figure 24. CAPA dose-response curves in HUVEC cells with or without OECs.** CAPA dose-response curves showing the degree of nuclear penetration for **(A)** ct-nusinersen and **(B)** ct-W in HUVEC cells in the presence or absence of 10 µM of UNC10217938A after 24 h incubation. All data show results from three independent trials and error bars show standard error of the mean. $CP_{50}$ values shown in **Figure 6C** and **SI Table 27**.

**SI Table 3. Sequences and molecular weights of the panel of compounds used in this study.** Notes: **(i)** For ct-SAHB, $S_5$ refers to *(S)*-2-(4′-pentenyl)alanine and the molecular weight reported is the one after stapling (see Methods). **(ii)** For ct-nusinersen, all the nucleotides had a 2'-methoxyethyl modification and, the backbone consisted of phosphorothioates and the compound was ordered with a 5' PS-azide linker (see **SI Figure 1**) from the Utah Core Facility. **(iii)** For ct-PMO, all the nucleotides had the PMO modification and the compound was ordered with a 5' azide from GeneTools. **(iv)** The chloroalkane carboxylic acid (RL-3180, Iris Biotech) linker was used for ct-R9W, ct-Tat, ct-SAHB, ct-W (see Methods) and the ct-DBCO (**SI Figure 1**) was used for ct-nusinersen and ct-PMO (see Methods).

| ct-compound | Amino acid/ nucleotide sequence | Expected molecular weight | Observed molecular weight |
|---|---|---|---|
| ct-R9W | RRRRRRRRRW | 1915.73 | 1915.47 |
| ct-Tat | YGRKKRRQRRR | 1864.67 | 1863.05 |
| ct-SAHB | IWIAQELR($S_5$)IGD($S_5$)FNAYYARR | 2910.85 | 2910.98 |
| ct-nusinersen | TCACTTTCATAATGCTGG | 7954.47 | 7954.58 |
| ct-PMO | GTTGCCTCCGGTTCTGAAGGTGTTC | 9288.12 | 9288.11 |
| ct-W | W | 509.04 | 509.32 |

**SI Table 4. Statistical differences in $CP_{50}$ values for ct-R9W in the seven cell lines.** (A) Summary of ordinary one-way ANOVA. (B) Summary of Tukey's multiple comparisons test.

**A.**

| **ANOVA summary** | |
|---|---|
| F | 44.18 |
| P value | <0.0001 |
| P value summary | **** |
| Significant diff. among means (P < 0.05)? | Yes |
| R squared | 0.9498 |

| **B. Tukey's multiple comparisons test** | **Mean Diff.** | **95.00% CI of diff.** | **Below threshold?** | **Summary** | **Adjusted P Value** |
|---|---|---|---|---|---|
| HeLa vs. Saos-2 | -0.03333 | -0.1696 to 0.1030 | No | ns | >0.9999 |
| HeLa vs. U2OS | 0.1358 | -0.0005035 to 0.2721 | No | ns | 0.0513 |
| HeLa vs. MIA-PaCa2 | 0.2870 | 0.1507 to 0.4233 | Yes | **** | <0.0001 |
| HeLa vs. HEK293T | -0.1135 | -0.2498 to 0.02277 | No | ns | 0.1605 |
| HeLa vs. BxPC3 | -0.08050 | -0.2168 to 0.05580 | No | ns | 0.6365 |
| HeLa vs. HepG2 | -0.2454 | -0.3817 to -0.1091 | Yes | *** | 0.0002 |
| Saos-2 vs. U2OS | 0.1691 | 0.03283 to 0.3054 | Yes | ** | 0.0090 |
| Saos-2 vs. MIA-PaCa2 | 0.3204 | 0.1841 to 0.4567 | Yes | **** | <0.0001 |
| Saos-2 vs. HEK293T | -0.08020 | -0.2165 to 0.05610 | No | ns | 0.6422 |
| Saos-2 vs. BxPC3 | -0.04717 | -0.1835 to 0.08914 | No | ns | 0.9950 |
| Saos-2 vs. HepG2 | -0.2120 | -0.3483 to -0.07573 | Yes | ** | 0.0011 |
| U2OS vs. MIA-PaCa2 | 0.1512 | 0.01492 to 0.2875 | Yes | * | 0.0229 |
| U2OS vs. HEK293T | -0.2493 | -0.3856 to -0.1130 | Yes | *** | 0.0002 |

| | | | | | |
|---|---|---|---|---|---|
| U2OS vs. BxPC3 | -0.2163 | -0.3526 to -0.08000 | Yes | *** | 0.0009 |
| U2OS vs. HepG2 | -0.3812 | -0.5175 to -0.2449 | Yes | **** | <0.0001 |
| MIA-PaCa2 vs. HEK293T | -0.4006 | -0.5369 to -0.2643 | Yes | **** | <0.0001 |
| MIA-PaCa2 vs. BxPC3 | -0.3675 | -0.5038 to -0.2312 | Yes | **** | <0.0001 |
| MIA-PaCa2 vs. HepG2 | -0.5324 | -0.6687 to -0.3961 | Yes | **** | <0.0001 |
| HEK293T vs. BxPC3 | 0.03303 | -0.1033 to 0.1693 | No | ns | >0.9999 |
| HEK293T vs. HepG2 | -0.1318 | -0.2681 to 0.004470 | No | ns | 0.0631 |
| BxPC3 vs. HepG2 | -0.1649 | -0.3012 to -0.02856 | Yes | * | 0.0113 |

**SI Table 5. Statistical differences in $CP_{50}$ values for ct-Tat in the seven cell lines.** (A) Summary of ordinary one-way ANOVA. (B) Summary of Tukey's multiple comparisons test.

**A.**

| **ANOVA summary** | |
|---|---|
| F | 39.75 |
| P value | <0.0001 |
| P value summary | **** |
| Significant diff. among means (P < 0.05)? | Yes |
| R squared | 0.9445 |

**B.**

| **Tukey's multiple comparisons test** | **Mean Diff.** | **95.00% CI of diff.** | **Below threshold?** | **Summary** | **Adjusted P Value** |
|---|---|---|---|---|---|
| HeLa vs. Saos-2 | 0.7400 | -0.8448 to 2.325 | No | ns | 0.6879 |
| HeLa vs. U2OS | 1.577 | -0.007826 to 3.162 | No | ns | 0.0515 |
| HeLa vs. MIA-PaCa2 | 2.734 | 1.149 to 4.318 | Yes | *** | 0.0006 |
| HeLa vs. HEK293T | -1.308 | -2.892 to 0.2772 | No | ns | 0.1397 |
| HeLa vs. BxPC3 | -3.356 | -4.941 to -1.772 | Yes | **** | <0.0001 |
| HeLa vs. HepG2 | -1.548 | -3.133 to 0.03683 | No | ns | 0.0576 |
| Saos-2 vs. U2OS | 0.8370 | -0.7478 to 2.422 | No | ns | 0.5664 |
| Saos-2 vs. MIA-PaCa2 | 1.994 | 0.4088 to 3.578 | Yes | * | 0.0101 |
| Saos-2 vs. HEK293T | -2.048 | -3.632 to -0.4628 | Yes | ** | 0.0082 |
| Saos-2 vs. BxPC3 | -4.096 | -5.681 to -2.512 | Yes | **** | <0.0001 |
| Saos-2 vs. HepG2 | -2.288 | -3.873 to -0.7032 | Yes | ** | 0.0032 |
| U2OS vs. MIA-PaCa2 | 1.157 | -0.4282 to 2.741 | No | ns | 0.2333 |
| U2OS vs. HEK293T | -2.885 | -4.469 to -1.300 | Yes | *** | 0.0004 |
| U2OS vs. BxPC3 | -4.933 | -6.518 to -3.349 | Yes | **** | <0.0001 |
| U2OS vs. HepG2 | -3.125 | -4.710 to -1.540 | Yes | *** | 0.0002 |
| MIA-PaCa2 vs. HEK293T | -4.041 | -5.626 to -2.456 | Yes | **** | <0.0001 |
| MIA-PaCa2 vs. BxPC3 | -6.090 | -7.675 to -4.505 | Yes | **** | <0.0001 |
| MIA-PaCa2 vs. HepG2 | -4.282 | -5.866 to -2.697 | Yes | **** | <0.0001 |
| HEK293T vs. BxPC3 | -2.049 | -3.633 to -0.4638 | Yes | ** | 0.0082 |
| HEK293T vs. HepG2 | -0.2403 | -1.825 to 1.344 | No | ns | 0.9981 |
| BxPC3 vs. HepG2 | 1.808 | 0.2235 to 3.393 | Yes | * | 0.0210 |

**SI Table 6. Statistical differences in $CP_{50}$ values for ct-SAHB in the seven cell lines.** (A) Summary of ordinary one-way ANOVA. (B) Summary of Tukey's multiple comparisons test.

**A.**

| **ANOVA summary** | |
|---|---|
| F | 19.04 |
| P value | <0.0001 |
| P value summary | **** |
| Significant diff. among means (P < 0.05)? | Yes |
| R squared | 0.8908 |

**B.**

| **Tukey's multiple comparisons test** | **Mean Diff.** | **95.00% CI of diff.** | **Below threshold?** | **Summary** | **Adjusted P Value** |
|---|---|---|---|---|---|
| HeLa vs. Saos-2 | 0.2942 | -0.2349 to 0.8233 | No | ns | 0.5115 |
| HeLa vs. U2OS | 0.3726 | -0.1565 to 0.9017 | No | ns | 0.2655 |
| HeLa vs. MIA-PaCa2 | 0.4112 | -0.1179 to 0.9403 | No | ns | 0.1819 |
| HeLa vs. HEK293T | -0.9154 | -1.445 to -0.3863 | Yes | *** | 0.0006 |
| HeLa vs. BxPC3 | -0.3584 | -0.8875 to 0.1707 | No | ns | 0.3027 |
| HeLa vs. HepG2 | 0.1067 | -0.4224 to 0.6358 | No | ns | 0.9912 |
| Saos-2 vs. U2OS | 0.07839 | -0.4507 to 0.6075 | No | ns | 0.9983 |
| Saos-2 vs. MIA-PaCa2 | 0.1169 | -0.4122 to 0.6460 | No | ns | 0.9859 |
| Saos-2 vs. HEK293T | -1.210 | -1.739 to -0.6806 | Yes | **** | <0.0001 |
| Saos-2 vs. BxPC3 | -0.6527 | -1.182 to -0.1236 | Yes | * | 0.0118 |
| Saos-2 vs. HepG2 | -0.1875 | -0.7166 to 0.3416 | No | ns | 0.8790 |
| U2OS vs. MIA-PaCa2 | 0.03854 | -0.4906 to 0.5677 | No | ns | >0.9999 |
| U2OS vs. HEK293T | -1.288 | -1.817 to -0.7589 | Yes | **** | <0.0001 |
| U2OS vs. BxPC3 | -0.7311 | -1.260 to -0.2019 | Yes | ** | 0.0047 |
| U2OS vs. HepG2 | -0.2659 | -0.7950 to 0.2632 | No | ns | 0.6173 |
| MIA-PaCa2 vs. HEK293T | -1.327 | -1.856 to -0.7975 | Yes | **** | <0.0001 |
| MIA-PaCa2 vs. BxPC3 | -0.7696 | -1.299 to -0.2405 | Yes | ** | 0.0030 |
| MIA-PaCa2 vs. HepG2 | -0.3045 | -0.8336 to 0.2246 | No | ns | 0.4745 |
| HEK293T vs. BxPC3 | 0.5570 | 0.02789 to 1.086 | Yes | * | 0.0362 |
| HEK293T vs. HepG2 | 1.022 | 0.4930 to 1.551 | Yes | *** | 0.0002 |
| BxPC3 vs. HepG2 | 0.4651 | -0.06398 to 0.9942 | No | ns | 0.1027 |

**SI Table 7. Statistical differences in $CP_{50}$ values for ct-W (4hours) in the seven cell lines.** (A) Summary of ordinary one-way ANOVA. (B) Summary of Tukey's multiple comparisons test.

**A.**

| **ANOVA summary** | |
|---|---|
| F | 31.62 |
| P value | <0.0001 |
| P value summary | **** |
| Significant diff. among means (P < 0.05)? | Yes |
| R squared | 0.9313 |

**B.**

| **Tukey's multiple comparisons test** | **Mean Diff.** | **95.00% CI of diff.** | **Below threshold?** | **Summary** | **Adjusted P Value** |
|---|---|---|---|---|---|
| HeLa vs. Saos-2 | -0.001667 | -0.01362 to 0.01029 | No | ns | 0.9988 |
| HeLa vs. U2OS | -0.02791 | -0.03987 to -0.01596 | Yes | **** | <0.0001 |
| HeLa vs. MIA-PaCa2 | 0.01034 | -0.001617 to 0.02229 | No | ns | 0.1115 |
| HeLa vs. HEK293T | -0.02515 | -0.03710 to -0.01320 | Yes | **** | <0.0001 |
| HeLa vs. BxPC3 | -0.003080 | -0.01503 to 0.008875 | No | ns | 0.9700 |
| HeLa vs. HepG2 | -0.01086 | -0.02281 to 0.001095 | No | ns | 0.0865 |
| Saos-2 vs. U2OS | -0.02625 | -0.03820 to -0.01429 | Yes | **** | <0.0001 |
| Saos-2 vs. MIA-PaCa2 | 0.01201 | 5.004e-005 to 0.02396 | Yes | * | 0.0487 |
| Saos-2 vs. HEK293T | -0.02348 | -0.03544 to -0.01153 | Yes | *** | 0.0002 |
| Saos-2 vs. BxPC3 | -0.001413 | -0.01337 to 0.01054 | No | ns | 0.9995 |
| Saos-2 vs. HepG2 | -0.009193 | -0.02115 to 0.002762 | No | ns | 0.1900 |
| U2OS vs. MIA-PaCa2 | 0.03825 | 0.02630 to 0.05021 | Yes | **** | <0.0001 |
| U2OS vs. HEK293T | 0.002763 | -0.009192 to 0.01472 | No | ns | 0.9823 |
| U2OS vs. BxPC3 | 0.02483 | 0.01288 to 0.03679 | Yes | **** | <0.0001 |
| U2OS vs. HepG2 | 0.01705 | 0.005098 to 0.02901 | Yes | ** | 0.0036 |
| MIA-PaCa2 vs. HEK293T | -0.03549 | -0.04744 to -0.02353 | Yes | **** | <0.0001 |
| MIA-PaCa2 vs. BxPC3 | -0.01342 | -0.02537 to -0.001463 | Yes | * | 0.0235 |
| MIA-PaCa2 vs. HepG2 | -0.02120 | -0.03315 to -0.009243 | Yes | *** | 0.0005 |
| HEK293T vs. BxPC3 | 0.02207 | 0.01012 to 0.03402 | Yes | *** | 0.0003 |
| HEK293T vs. HepG2 | 0.01429 | 0.002335 to 0.02624 | Yes | * | 0.0150 |
| BxPC3 vs. HepG2 | -0.007780 | -0.01973 to 0.004175 | No | ns | 0.3432 |

**SI Table 8. Statistical differences in $CP_{50}$ values for ct-nusinersen in the seven cell lines.** (A) Summary of ordinary one-way ANOVA. (B) Summary of Tukey's multiple comparisons test.

**A.**

| ANOVA summary | |
|---|---|
| F | 13.86 |
| P value | <0.0001 |
| P value summary | **** |
| Significant diff. among means (P < 0.05)? | Yes |
| R squared | 0.8559 |

**B.**

| Tukey's multiple comparisons test | Mean Diff. | 95.00% CI of diff. | Below threshold? | Summary | Adjusted P Value |
|---|---|---|---|---|---|
| HeLa vs. Saos-2 | 0.2767 | 0.01362 to 0.5397 | Yes | * | 0.0364 |
| HeLa vs. U2OS | -0.2479 | -0.5109 to 0.01514 | No | ns | 0.0707 |
| HeLa vs. MIA-PaCa2 | 0.3427 | 0.07970 to 0.6058 | Yes | ** | 0.0077 |
| HeLa vs. HEK293T | -0.02663 | -0.2897 to 0.2364 | No | ns | 0.9998 |
| HeLa vs. BxPC3 | 0.05003 | -0.2130 to 0.3131 | No | ns | 0.9935 |
| HeLa vs. HepG2 | -0.06200 | -0.3250 to 0.2010 | No | ns | 0.9805 |
| Saos-2 vs. U2OS | -0.5246 | -0.7876 to -0.2615 | Yes | *** | 0.0001 |
| Saos-2 vs. MIA-PaCa2 | 0.06608 | -0.1970 to 0.3291 | No | ns | 0.9734 |
| Saos-2 vs. HEK293T | -0.3033 | -0.5663 to -0.04026 | Yes | * | 0.0195 |
| Saos-2 vs. BxPC3 | -0.2266 | -0.4897 to 0.03641 | No | ns | 0.1136 |
| Saos-2 vs. HepG2 | -0.3387 | -0.6017 to -0.07562 | Yes | ** | 0.0084 |
| U2OS vs. MIA-PaCa2 | 0.5906 | 0.3276 to 0.8537 | Yes | **** | <0.0001 |
| U2OS vs. HEK293T | 0.2213 | -0.04178 to 0.4843 | No | ns | 0.1276 |
| U2OS vs. BxPC3 | 0.2979 | 0.03489 to 0.5610 | Yes | * | 0.0221 |
| U2OS vs. HepG2 | 0.1859 | -0.07714 to 0.4489 | No | ns | 0.2622 |
| MIA-PaCa2 vs. HEK293T | -0.3694 | -0.6324 to -0.1063 | Yes | ** | 0.0041 |
| MIA-PaCa2 vs. BxPC3 | -0.2927 | -0.5558 to -0.02967 | Yes | * | 0.0250 |
| MIA-PaCa2 vs. HepG2 | -0.4047 | -0.6678 to -0.1417 | Yes | ** | 0.0018 |
| HEK293T vs. BxPC3 | 0.07667 | -0.1864 to 0.3397 | No | ns | 0.9469 |
| HEK293T vs. HepG2 | -0.03537 | -0.2984 to 0.2277 | No | ns | 0.9990 |
| BxPC3 vs. HepG2 | -0.1120 | -0.3751 to 0.1510 | No | ns | 0.7653 |

**SI Table 9. Statistical differences in $CP_{50}$ values for <u>ct-PMO</u> in the seven cell lines.** (A) Summary of ordinary one-way ANOVA. (B) Summary of Tukey's multiple comparisons test.

**A.**

| **ANOVA summary** | |
|---|---|
| F | 10.94 |
| P value | 0.0001 |
| P value summary | *** |
| Significant diff. among means (P < 0.05)? | Yes |
| R squared | 0.8242 |

**B.**

| **Tukey's multiple comparisons test** | **Mean Diff.** | **95.00% CI of diff.** | **Below threshold?** | **Summary** | **Adjusted P Value** |
|---|---|---|---|---|---|
| HeLa vs. Saos-2 | 0.04000 | -0.07241 to 0.1524 | No | ns | 0.8771 |
| HeLa vs. U2OS | 0.01166 | -0.1007 to 0.1241 | No | ns | 0.9998 |
| HeLa vs. MIA-PaCa2 | 0.03376 | -0.07865 to 0.1462 | No | ns | 0.9393 |
| HeLa vs. HEK293T | -0.1767 | -0.2891 to -0.06426 | Yes | ** | 0.0015 |
| HeLa vs. BxPC3 | 0.04309 | -0.06932 to 0.1555 | No | ns | 0.8372 |
| HeLa vs. HepG2 | -0.007600 | -0.1200 to 0.1048 | No | ns | >0.9999 |
| Saos-2 vs. U2OS | -0.02834 | -0.1407 to 0.08407 | No | ns | 0.9730 |
| Saos-2 vs. MIA-PaCa2 | -0.006237 | -0.1186 to 0.1062 | No | ns | >0.9999 |
| Saos-2 vs. HEK293T | -0.2167 | -0.3291 to -0.1043 | Yes | *** | 0.0002 |
| Saos-2 vs. BxPC3 | 0.003090 | -0.1093 to 0.1155 | No | ns | >0.9999 |
| Saos-2 vs. HepG2 | -0.04760 | -0.1600 to 0.06481 | No | ns | 0.7698 |
| U2OS vs. MIA-PaCa2 | 0.02210 | -0.09031 to 0.1345 | No | ns | 0.9923 |
| U2OS vs. HEK293T | -0.1883 | -0.3007 to -0.07592 | Yes | *** | 0.0008 |
| U2OS vs. BxPC3 | 0.03143 | -0.08098 to 0.1438 | No | ns | 0.9560 |
| U2OS vs. HepG2 | -0.01926 | -0.1317 to 0.09315 | No | ns | 0.9963 |
| MIA-PaCa2 vs. HEK293T | -0.2104 | -0.3228 to -0.09802 | Yes | *** | 0.0003 |
| MIA-PaCa2 vs. BxPC3 | 0.009327 | -0.1031 to 0.1217 | No | ns | >0.9999 |
| MIA-PaCa2 vs. HepG2 | -0.04136 | -0.1538 to 0.07105 | No | ns | 0.8602 |
| HEK293T vs. BxPC3 | 0.2198 | 0.1073 to 0.3322 | Yes | *** | 0.0002 |
| HEK293T vs. HepG2 | 0.1691 | 0.05666 to 0.2815 | Yes | ** | 0.0022 |
| BxPC3 vs. HepG2 | -0.05069 | -0.1631 to 0.06172 | No | ns | 0.7188 |

**SI Table 10. Statistical differences in $CP_{50}$ values for ct-R9W (24hours) in the seven cell lines.** (A) Summary of ordinary one-way ANOVA. (B) Summary of Tukey's multiple comparisons test.

**A.**

| **ANOVA summary** | |
|---|---|
| F | 5.530 |
| P value | 0.0040 |
| P value summary | ** |
| Significant diff. among means (P < 0.05)? | Yes |
| R squared | 0.7032 |

**B.**

| **Tukey's multiple comparisons test** | **Mean Diff.** | **95.00% CI of diff.** | **Below threshold?** | **Summary** | **Adjusted P Value** |
|---|---|---|---|---|---|
| HeLa vs. Saos-2 | -0.09333 | -0.2587 to 0.07204 | No | ns | 0.4955 |
| HeLa vs. U2OS | -0.0006333 | -0.1660 to 0.1647 | No | ns | >0.9999 |
| HeLa vs. MIA-PaCa2 | 0.1075 | -0.05790 to 0.2729 | No | ns | 0.3446 |
| HeLa vs. HEK293T | -0.05840 | -0.2238 to 0.1070 | No | ns | 0.8807 |
| HeLa vs. BxPC3 | -0.06447 | -0.2298 to 0.1009 | No | ns | 0.8269 |
| HeLa vs. HepG2 | -0.1417 | -0.3071 to 0.02364 | No | ns | 0.1166 |
| Saos-2 vs. U2OS | 0.09270 | -0.07267 to 0.2581 | No | ns | 0.5029 |
| Saos-2 vs. MIA-PaCa2 | 0.2008 | 0.03544 to 0.3662 | Yes | * | 0.0133 |
| Saos-2 vs. HEK293T | 0.03493 | -0.1304 to 0.2003 | No | ns | 0.9888 |
| Saos-2 vs. BxPC3 | 0.02887 | -0.1365 to 0.1942 | No | ns | 0.9959 |
| Saos-2 vs. HepG2 | -0.04840 | -0.2138 to 0.1170 | No | ns | 0.9459 |
| U2OS vs. MIA-PaCa2 | 0.1081 | -0.05726 to 0.2735 | No | ns | 0.3385 |
| U2OS vs. HEK293T | -0.05777 | -0.2231 to 0.1076 | No | ns | 0.8858 |
| U2OS vs. BxPC3 | -0.06383 | -0.2292 to 0.1015 | No | ns | 0.8330 |
| U2OS vs. HepG2 | -0.1411 | -0.3065 to 0.02427 | No | ns | 0.1192 |
| MIA-PaCa2 vs. HEK293T | -0.1659 | -0.3313 to -0.0005017 | Yes | * | 0.0491 |
| MIA-PaCa2 vs. BxPC3 | -0.1719 | -0.3373 to -0.006568 | Yes | * | 0.0392 |
| MIA-PaCa2 vs. HepG2 | -0.2492 | -0.4146 to -0.08384 | Yes | ** | 0.0022 |
| HEK293T vs. BxPC3 | -0.006067 | -0.1714 to 0.1593 | No | ns | >0.9999 |
| HEK293T vs. HepG2 | -0.08333 | -0.2487 to 0.08204 | No | ns | 0.6147 |
| BxPC3 vs. HepG2 | -0.07727 | -0.2426 to 0.08811 | No | ns | 0.6874 |

**SI Table 11. Statistical differences in $CP_{50}$ values for ct-W (24hours) in the seven cell lines.** (A) Summary of ordinary one-way ANOVA. (B) Summary of Tukey's multiple comparisons test.

**A.**

| **ANOVA summary** | |
|---|---|
| F | 46.25 |
| P value | <0.0001 |
| P value summary | **** |
| Significant diff. among means (P < 0.05)? | Yes |
| R squared | 0.9520 |

**B.**

| **Tukey's multiple comparisons test** | **Mean Diff.** | **95.00% CI of diff.** | **Below threshold?** | **Summary** | **Adjusted P Value** |
|---|---|---|---|---|---|
| HeLa vs. Saos-2 | 3.333e-005 | -0.002075 to 0.002142 | No | ns | >0.9999 |
| HeLa vs. U2OS | -0.006686 | -0.008794 to -0.004577 | Yes | **** | <0.0001 |
| HeLa vs. MIA-PaCa2 | 0.001636 | -0.0004723 to 0.003744 | No | ns | 0.1830 |
| HeLa vs. HEK293T | -0.002566 | -0.004674 to -0.0004578 | Yes | * | 0.0131 |
| HeLa vs. BxPC3 | 0.0009370 | -0.001171 to 0.003045 | No | ns | 0.7311 |
| HeLa vs. HepG2 | -0.003620 | -0.005728 to -0.001512 | Yes | *** | 0.0006 |
| Saos-2 vs. U2OS | -0.006719 | -0.008827 to -0.004611 | Yes | **** | <0.0001 |
| Saos-2 vs. MIA-PaCa2 | 0.001603 | -0.0005056 to 0.003711 | No | ns | 0.1991 |
| Saos-2 vs. HEK293T | -0.002599 | -0.004708 to -0.0004911 | Yes | * | 0.0118 |
| Saos-2 vs. BxPC3 | 0.0009037 | -0.001205 to 0.003012 | No | ns | 0.7604 |
| Saos-2 vs. HepG2 | -0.003653 | -0.005762 to -0.001545 | Yes | *** | 0.0006 |
| U2OS vs. MIA-PaCa2 | 0.008322 | 0.006213 to 0.01043 | Yes | **** | <0.0001 |
| U2OS vs. HEK293T | 0.004120 | 0.002011 to 0.006228 | Yes | *** | 0.0002 |
| U2OS vs. BxPC3 | 0.007623 | 0.005514 to 0.009731 | Yes | **** | <0.0001 |
| U2OS vs. HepG2 | 0.003066 | 0.0009575 to 0.005174 | Yes | ** | 0.0030 |
| MIA-PaCa2 vs. HEK293T | -0.004202 | -0.006310 to -0.002094 | Yes | *** | 0.0001 |
| MIA-PaCa2 vs. BxPC3 | -0.0006989 | -0.002807 to 0.001409 | No | ns | 0.9075 |
| MIA-PaCa2 vs. HepG2 | -0.005256 | -0.007364 to -0.003148 | Yes | **** | <0.0001 |
| HEK293T vs. BxPC3 | 0.003503 | 0.001395 to 0.005611 | Yes | *** | 0.0009 |
| HEK293T vs. HepG2 | -0.001054 | -0.003162 to 0.001054 | No | ns | 0.6226 |
| BxPC3 vs. HepG2 | -0.004557 | -0.006665 to -0.002449 | Yes | **** | <0.0001 |

**SI Table 12. $CP_{50}$ values for ct-compounds in the seven cell lines (using the narrow gate).** $CP_{50}$ values are reported as the average standard error of the mean from three independent curve fits to three independent trials.

| Cell Line | 4 hours | | | | 24 hours | | | |
|---|---|---|---|---|---|---|---|---|
| | ct-R9W | ct-Tat | ct-SAHB | ct-W | ct-nusinersen | ct-PMO | ct-R9W | ct-W |
| **HeLa** | 0.36 ± 0.02 | 3.68 ± 0.48 | 0.56 ± 0.12 | 0.014 ± 0.001 | 0.42 ± 0.09 | 0.15 ± 0.02 | 0.13 ± 0.02 | 0.003 ± 0.0006 |
| **Saos-2** | 0.38 ± 0.04 | 2.94 ± 0.26 | 0.27 ± 0.06 | 0.015 ± 0.002 | 0.15 ± 0.03 | 0.11 ± 0.03 | 0.23 ± 0.02 | 0.003 ± 0.0003 |
| **U2OS** | 0.22 ± 0.02 | 2.10 ± 0.15 | 0.19 ± 0.05 | 0.04 ± 0.004 | 0.67 ± 0.03 | 0.14 ± 0.03 | 0.13 ± 0.003 | 0.009 ± 0.0003 |
| **MIA PaCa-2** | 0.07 ± 0.01 | 0.94 ± 0.09 | 0.15 ± 0.01 | 0.003 ± 0.0002 | 0.08 ± 0.01 | 0.11 ± 0.02 | 0.03 ± 0.01 | 0.001 ± 0.0002 |
| **HEK293T** | 0.47 ± 0.01 | 4.98 ± 0.26 | 1.48 ± 0.13 | 0.04 ± 0.004 | 0.45 ± 0.05 | 0.32 ± 0.02 | 0.19 ± 0.03 | 0.005 ± 0.001 |
| **BxPC3** | 0.43 ± 0.03 | 7.03 ± 0.57 | 0.92 ± 0.17 | 0.02 ± 0.002 | 0.37 ± 0.04 | 0.10 ± 0.02 | 0.20 ± 0.01 | 0.002 ± 0.0003 |
| **HepG2** | 0.60 ± 0.04 | 5.22 ± 0.18 | 0.46 ± 0.13 | 0.02 ± 0.002 | 0.49 ± 0.10 | 0.15 ± 0.03 | 0.28 ± 0.08 | 0.006 ± 0.001 |

**SI Table 13. $CP_{50}$ values for ct-compounds in the seven cell lines (using the wide gate).** $CP_{50}$ values are reported as the average standard error of the mean from three independent curve fits to three independent trials.

| Cell Line | 4 hours | | | | 24 hours | | | |
|---|---|---|---|---|---|---|---|---|
| | ct-R9W | ct-Tat | ct-SAHB | ct-W | ct-nusinersen | ct-PMO | ct-R9W | ct-W |
| **HeLa** | 0.39 ± 0.03 | 5.32 ± 0.91 | 1.01 ± 0.23 | 0.026 ± 0.01 | 1.24 ± 0.41 | 0.32 ± 0.13 | 0.18 ± 0.04 | 0.008± 0.005 |
| **Saos-2** | 0.45 ± 0.03 | 3.80 ± 0.37 | 0.53 ± 0.08 | 0.021 ± 0.002 | 0.27 ± 0.08 | 0.17 ± 0.07 | 0.29 ± 0.08 | 0.003 ± 0.001 |
| **U2OS** | 0.24 ± 0.02 | 3.06 ± 0.28 | 0.49 ± 0.19 | 0.066 ± 0.009 | 1.09 ± 0.06 | 0.22 ± 0.07 | 0.15 ± 0.01 | 0.015 ± 0.001 |
| **MIA PaCa-2** | 0.067 ± 0.011 | 1.37 ± 0.14 | 0.28 ± 0.03 | 0.0044 ± 0.0004 | 0.12 ± 0.02 | 0.15 ± 0.03 | 0.03 ± 0.01 | 0.001 ± 0.0003 |
| **HEK293T** | 0.58 ± 0.02 | 6.36 ± 1.45 | 3.11 ± 0.72 | 0.06 ± 0.01 | 0.78 ± 0.05 | 0.68 ± 0.12 | 0.24 ± 0.05 | 0.010 ± 0.003 |
| **BxPC3** | 0.40 ± 0.07 | 8.02 ± 0.19 | 0.88 ± 0.20 | 0.02 ± 0.004 | 0.61 ± 0.10 | 0.17 ± 0.02 | 0.27 ± 0.02 | 0.004 ± 0.0003 |
| **HepG2** | 0.61 ± 0.04 | 6.81 ± 0.50 | 0.70 ± 0.20 | 0.03 ± 0.003 | 0.63 ± 0.09 | 0.23 ± 0.03 | 0.34 ± 0.11 | 0.010 ± 0.001 |

**SI Table 14. Relative fluorescence values of endocytosis markers in the seven cell lines.** Cells were treated with 1mg/mL TAMRA-dextran (high molecular weight), 1mg/mL TAMRA-transferrin or 5µg/mL AF647-CTB for 1 hour at 37°C, washed for 30mins, and then fluorescence was measured using benchtop flow cytometry.

| Cell Line | Dextran | Transferrin | Cholera Toxin subunit B |
|---|---|---|---|
| **HeLa** | 2.84 ± 0.15 | 4.68 ± 0.12 | 20.62 ± 0.74 |
| **Saos-2** | 3.81 ± 0.20 | 4.06 ± 0.16 | 20.01 ± 0.20 |
| **U2OS** | 2.84 ± 0.06 | 5.23 ± 0.02 | 14.88 ± 0.51 |
| **MIA PaCa-2** | 3.63 ± 0.17 | 5.5 ± 0.33 | 27.44 ± 3.49 |
| **HEK293T** | 9.77 ± 0.62 | 10.82 ± 0.17 | 66.64 ± 4.13 |
| **BxPC3** | 2.58 ± 0.10 | 3.16 ± 0.04 | 37.69 ± 8.40 |
| **HepG2** | 1.99 ± 0.05 | 2.54 ± 0.08 | 19.95 ± 1.83 |

**SI Table 15. CAPA $CP_{50}$ values in BxPC3 cells with or without chloroquine.**

| | ct-compound | No CQ | 60μM CQ |
|---|---|---|---|
| **4 hours** | ct-R9W | 0.40 ± 0.07 | 0.59 ± 0.12 |
| | ct-Tat | 8.02 ± 0.19 | 9.66 ± 1.69 |
| | ct-SAHB | 0.88 ± 0.20 | 6.44 ± 0.17 |
| | ct-W | 0.02 ± 0.004 | 0.03 ± 0.005 |
| **24 hours** | ct-nusinersen | 0.61 ± 0.1 | 1.97 ± 0.17 |
| | ct-PMO | 0.17 ± 0.02 | 0.43 ± 0.09 |
| | ct-R9W | 0.27 ± 0.02 | 0.58 ± 0.16 |
| | ct-W | 0.004 ± 0.0003 | 0.008 ± 0.001 |

**SI Table 16. CAPA $CP_{50}$ values in BxPC3 cells with or without OECs.** $CP_{50}$ values (μM) for ct-nusinersen, ct-PMO, and ct-W in the absence of OEC or in the presence of 10 μM UNC10217938A or 5 μM SH-BC-893 in **BxPC3 cells**.

| ct-compound | No OEC | UNC10217938A | SH-BC-893 |
|---|---|---|---|
| ct-nusinersen | 0.61 ± 0.1 | 0.43 ± 0.06 | 0.39 ± 0.08 |
| ct-PMO | 0.17 ± 0.02 | 0.21 ± 0.02 | 0.22 ± 0.04 |
| ct-W | 0.004 ± 0.0003 | 0.006 ± 0.0003 | 0.005 ± 0.0001 |

**SI Table 17. CAPA $CP_{50}$ values in U2OS cells with or without OECs.** $CP_{50}$ values (μM) for ct-nusinersen, ct-PMO, and ct-W in the absence of OECs or in the presence of 10μM UNC10217938A or 5μM SH-BC-893 in **U2OS cells**.

| ct-compound | No OEC | UNC10217938A | SH-BC-893 |
|---|---|---|---|
| ct-nusinersen | 0.50 ± 0.02 | 0.17 ± 0.08 | 0.48 ± 0.01 |
| ct-PMO | 0.33 ± 0.03 | 0.33 ± 0.01 | 0.29 ± 0.02 |
| ct-W | 0.009 ± 0.0003 | 0.003 ± 0.0002 | 0.007 ± 0.001 |

**SI Table 18. Statistical differences in $CP_{50}$ values for <u>ct-nusinersen (24hours)</u> in the BxPC3 cells with or without OECs.** (A) Summary of ordinary one-way ANOVA. (B) Summary of Tukey's multiple comparisons test.

**A.**

| ANOVA summary | |
|---|---|
| F | 1.922 |
| P value | 0.2264 |
| P value summary | ns |
| Significant diff. among means (P < 0.05)? | No |
| R squared | 0.3905 |

**B.**

| Tukey's multiple comparisons test | Mean Diff. | 95.00% CI of diff. | Below threshold? | Summary | Adjusted P Value |
|---|---|---|---|---|---|
| ct-nusinersen vs. ct-nusinersen + UNC10217938A | 0.1757 | -0.1802 to 0.5316 | No | ns | 0.3495 |
| ct-nusinersen vs. ct-nusinersen + SH-BC-893 | 0.2129 | -0.1430 to 0.5689 | No | ns | 0.2371 |
| ct-nusinersen + UNC10217938A vs. ct-nusinersen + SH-BC-893 | 0.03723 | -0.3187 to 0.3932 | No | ns | 0.9453 |

**SI Table 19. Statistical differences in $CP_{50}$ values for <u>ct-PMO (24hours)</u> in the BxPC3 cells with or without OECs.** (A) Summary of ordinary one-way ANOVA. (B) Summary of Tukey's multiple comparisons test.

**A.**

| ANOVA summary | |
|---|---|
| F | 1.099 |
| P value | 0.3921 |
| P value summary | ns |
| Significant diff. among means (P < 0.05)? | No |
| R squared | 0.2681 |

**B.**

| Tukey's multiple comparisons test | Mean Diff. | 95.00% CI of diff. | Below threshold? | Summary | Adjusted P Value |
|---|---|---|---|---|---|
| ct-PMO vs. ct-PMO + UNC10217938A | -0.04283 | -0.1554 to 0.06978 | No | ns | 0.5125 |
| ct-PMO vs. ct-PMO + SH-BC-893 | -0.05047 | -0.1631 to 0.06215 | No | ns | 0.4100 |
| ct-PMO + UNC10217938A vs. ct-PMO + SH-BC-893 | -0.007633 | -0.1202 to 0.1050 | No | ns | 0.9765 |

**SI Table 20. Statistical differences in $CP_{50}$ values for ct-nusinersen (24hours) in the U2OS cells with or without OECs.** (A) Summary of ordinary one-way ANOVA. (B) Summary of Tukey's multiple comparisons test.

**A.**

| ANOVA summary | |
|---|---|
| F | 14.97 |
| P value | 0.0047 |
| P value summary | ** |
| Significant diff. among means (P < 0.05)? | Yes |
| R squared | 0.8331 |

**B.**

| Tukey's multiple comparisons test | Mean Diff. | 95.00% CI of diff. | Below threshold? | Summary | Adjusted P Value |
|---|---|---|---|---|---|
| ct-nusinersen vs. ct-nusinersen + UNC10217938A | 0.3339 | 0.1257 to 0.5420 | Yes | ** | 0.0063 |
| ct-nusinersen vs. ct-nusinersen + SH-BC-893 | 0.02640 | -0.1818 to 0.2346 | No | ns | 0.9210 |
| ct-nusinersen + UNC10217938A vs. ct-nusinersen + SH-BC-893 | -0.3075 | -0.5156 to -0.09934 | Yes | ** | 0.0094 |

**SI Table 21. Statistical differences in $CP_{50}$ values for ct-PMO (24hours) in the U2OS cells with or without OECs.** (A) Summary of ordinary one-way ANOVA. (B) Summary of Tukey's multiple comparisons test.

**A.**

| ANOVA summary | |
|---|---|
| F | 1.528 |
| P value | 0.2908 |
| P value summary | ns |
| Significant diff. among means (P < 0.05)? | No |
| R squared | 0.3375 |

**B.**

| Tukey's multiple comparisons test | Mean Diff. | 95.00% CI of diff. | Below threshold? | Summary | Adjusted P Value |
|---|---|---|---|---|---|
| ct-PMO vs. ct-PMO + UNC10217938A | 0.002900 | -0.08799 to 0.09379 | No | ns | 0.9947 |
| ct-PMO vs. ct-PMO + SH-BC-893 | 0.04623 | -0.04466 to 0.1371 | No | ns | 0.3310 |
| ct-PMO + UNC10217938A vs. ct-PMO + SH-BC-893 | 0.04333 | -0.04756 to 0.1342 | No | ns | 0.3711 |

**SI Table 22. Statistical differences in $CP_{50}$ values for ct-W (24hours) in the BxPC3 cells with or without OECs.** (A) Summary of ordinary one-way ANOVA. (B) Summary of Tukey's multiple comparisons test.

**A.**

| **ANOVA summary** | |
|---|---|
| F | 25.60 |
| P value | 0.0012 |
| P value summary | ** |
| Significant diff. among means (P < 0.05)? | Yes |
| R squared | 0.8951 |

**B.**

| Tukey's multiple comparisons test | Mean Diff. | 95.00% CI of diff. | Below threshold? | Summary | Adjusted P Value |
|---|---|---|---|---|---|
| ct-W vs. ct-W + UNC10217938A | -0.002516 | -0.003599 to -0.001432 | Yes | *** | 0.0009 |
| ct-W vs. ct-W + SH-BC-893 | -0.001462 | -0.002546 to -0.0003785 | Yes | * | 0.0143 |
| ct-W + UNC10217938A vs. ct-W + SH-BC-893 | 0.001054 | -2.985e-005 to 0.002137 | No | ns | 0.0555 |

**SI Table 23: Statistical differences in $CP_{50}$ values for ct-W (24hours) in the U2OS cells with or without OECs.** (A) Summary of ordinary one-way ANOVA. (B) Summary of Tukey's multiple comparisons test.

**A.**

| **ANOVA summary** | |
|---|---|
| F | 31.05 |
| P value | 0.0007 |
| P value summary | *** |
| Significant diff. among means (P < 0.05)? | Yes |
| R squared | 0.9119 |

**B.**

| Tukey's multiple comparisons test | Mean Diff. | 95.00% CI of diff. | Below threshold? | Summary | Adjusted P Value |
|---|---|---|---|---|---|
| ct-W vs. ct-W + UNC10217938A | 0.005992 | 0.003630 to 0.008354 | Yes | *** | 0.0006 |
| ct-W vs. ct-W + SH-BC-893 | 0.002172 | -0.0001902 to 0.004534 | No | ns | 0.0680 |
| ct-W + UNC10217938A vs. ct-W + SH-BC-893 | -0.003820 | -0.006182 to -0.001458 | Yes | ** | 0.0061 |

**SI Table 24. CAPA $CP_{50}$ values in MIA PaCa-2 before or after clathrin knockdown.**

| | ct-compound | MIA PaCa-2 (WT) | Clathrin knockdown |
|---|---|---|---|
| 4 hours | ct-R9W | 0.067 ± 0.01 | 0.22 ± 0.04 |
| | ct-Tat | 1.37 ± 0.14 | 3.83±0.36 |
| | ct-SAHB | 0.28 ± 0.03 | 1.50 ± 0.08 |
| | ct-W | 0.004 ± 0.0004 | 0.011 ± 0.003 |
| 24 hours | ct-nusinersen | 0.12 ± 0.02 | 0.22 ± 0.05 |
| | ct-PMO | 0.15 ± 0.03 | 0.18 ± 0.03 |
| | ct-R9W | 0.03 ± 0.01 | 0.06 ± 0.01 |
| | ct-W | 0.0015 ± 0.0003 | 0.0016 ± 0.0007 |

**SI Table 25. Unpaired t-test comparing the markers uptake before and after clathrin knockdown in MIA-PaCa-2 cells.**

| Unpaired t test | Dextran uptake | Transferrin uptake | CTB uptake |
|---|---|---|---|
| P value | 0.0615 | 0.0801 | <0.0001 |
| P value summary | ns | ns | **** |
| Significantly different (P < 0.05)? | No | No | Yes |
| One- or two-tailed P value? | Two-tailed | Two-tailed | Two-tailed |
| t, df | t=2.295, df=6 | t=2.104, df=6 | t=9.182, df=6 |

**SI Table 26. Unpaired t-test comparing the $CP_{50}$s of ct-compounds before and after clathrin knockdown in MIA-PaCa-2 cells.**

| Unpaired t test | ct-R9W, 4 h | ct-Tat, 4 h | ct-SAHB, 4 h | ct-W, 4 h |
|---|---|---|---|---|
| P value | 0.0137 | 0.0030 | 0.0001 | 0.0860 |
| P value summary | * | ** | *** | ns |
| Significantly different (P < 0.05)? | Yes | Yes | Yes | No |
| One- or two-tailed P value? | Two-tailed | Two-tailed | Two-tailed | Two-tailed |
| t, df | t=4.196, df=4 | t=6.409, df=4 | t=14.27, df=4 | t=2.267, df=4 |

| Unpaired t test | ct-nusinersen, 24 h | ct-PMO, 4 h | ct-R9W, 24 h | ct-W, 24 h |
|---|---|---|---|---|
| P value | 0.1114 | 0.4720 | 0.1334 | 0.9080 |
| P value summary | ns | ns | ns | ns |
| Significantly different (P < 0.05)? | No | No | No | No |
| One- or two-tailed P value? | Two-tailed | Two-tailed | Two-tailed | Two-tailed |
| t, df | t=2.036, df=4 | t=0.7934, df=4 | t=1.879, df=4 | t=0.1231, df=4 |

**SI Table 27. CAPA $CP_{50}$ values in HUVEC cells with or without OECs.** $CP_{50}$ values are reported as the average and standard error of the mean from three independent curve fits to three independent trials.

| ct-compound | NO OEC | +10μM UNC10217938A |
|---|---|---|
| ct-nusinersen | 1.59 ± 0.28 | 0.94 ± 0.12 |
| ct-W | 0.0024 ± 0.0003 | 0.0030 ± 0.0001 |
